# Single-molecule chromosome tracing reveals a diversity of megabase heterochromatin domains

**DOI:** 10.1101/2025.09.03.673923

**Authors:** Dania Camila Pulido Barrera, Jia E. You, Fei Xu, Tobias Verheijen, Ahilya N. Sawh, Pavel Kos, Luca Giorgetti, Nacho Molina, David Brückner, Susan E. Mango

## Abstract

Chromosomes fold into distinct domains that regulate transcription, replication, and repair. Beyond well-characterized TADs and compartments, the diversity of heterochromatin domains remains poorly defined at the sequence level. Using single-molecule tracing of nascent heterochromatin in *C. elegans*, we identify three classes of megabase-scale domains: (i) sharp-boundary, Condensin I–dependent Topological Associating Domain-like domains (TADLs); (ii) similarly sized, but Condensin-independent, *<u>el</u>egans* <u>Condensin</u>-Independent <u>D</u>omains (elCIDs); and (iii) weaker, diffuse structures that are abundant in the population. TADLs arise early in development, preceding elCIDs, and both become progressively compacted through H3K9 methylation, which promotes intra- and inter-domain proximity. Condensin mutations disrupt TADLs, yet single molecules can still form domain-like structures, as recapitulated by free polymer simulations. However, these differ markedly in boundary positioning and biophysical properties. Our results uncover previously unrecognized heterochromatin architectures and demonstrate that single-molecule analysis and mutational dissection provide valuable approaches for distinguishing between domain types.

## Introduction

Recent advances in chromatin biology have shown that the genome is organized into domains across multiple scales. At the megabase scale, Hi-C studies revealed topologically associating domains (TADs), while specific histone modifications contribute to the segregation of chromatin into euchromatic and heterochromatic compartments ^1–4^. These structures play key roles in gene regulation, genome integrity, and higher-order nuclear organization ^5–9^ Beyond these Hi-C-defined domains, microscopy-based approaches have revealed additional physical structures such as packing domains (PDs), Polycomb bodies, and the nucleolus, which reflect chromatin density and organization at diverse scales ^10–15^. These structures are distinct from TADs. PDs, for instance, do not always coincide with Hi-C–defined TADs or compartments, suggesting distinct and complementary modes of genome folding that elude crosslinking-based approaches.

Early models proposed a hierarchical architecture of multi-megabase A (euchromatic) and B (heterochromatic) compartments subdivided into TADs ^3,16,17^. High-resolution 3C-based maps now reveal a more dynamic, multilayered configuration, where CTCF-anchored loops, compartments, and finer-scale features coexist ^18,19^. These insights highlight the limitations of defining chromatin domains by size or contact pattern alone, and emphasize the need for mechanistic classifications. In higher organisms, loop extrusion by SMC complexes plays a central role in shaping chromatin domains. Cohesin drives transient enhancer-promoter loops within TADs ^7,20–27^. Condensin, another SMC complex, also builds chromatin domains in interphase, as shown in vertebrate cells and *C. elegans* ^28–30^. Packing domains (PDs) are thought to form through the combined action of transcription, nucleosome remodeling, and loop extrusion, but once stabilized, they persist independently of the Cohesin subunit Rad21 ^10,11^, underscoring how multiple molecular processes work together to shape genome organization.

In *C. elegans*, megabase-scale TADs have been observed for the X chromosome, where a specialized Condensin I complex (Condensin I^DC^) and its associated proteins generate both megabase-sized TADs and TAD boundaries ^23,31^. Depletion of Condensin complexes disrupts TADs on the X chromosome and interactions across both autosomes and the X chromosome, leading to reduced contacts ranging from 100 bp to 2 Mb ^24^. This disruption suggests that loop extrusion influences chromatin organization on autosomes, however clear TADs with strong boundaries have not been seen at the megabase scale ^23,24,32^. Instead, autosomes feature small TAD-like domains, each spanning a few genes as in vertebrates but with sizes ranging from 20 to 40 kb ^33^. Small loops rely on Cohesin for their formation, similar to the larger TADs in mammals ^34,35^. These observations suggest that Cohesin in *C. elegans* generates small TADL domains, and begs the question of whether larger looped domains exist, similar to those in vertebrates.

In this study, we explore the organization of autosomes at the megabase scale. We focus on a stretch of heterochromatin that is known to undergo large-scale folding ^36^. Using chromosome tracing to achieve single-chromosome resolution ^36–38^, we describe chromatin domains at the megabase scale, with variable boundary strengths and genetic requirements. Some resemble TADs, with sharp boundaries and a need for loop extrusion by Condensin I, while others are independent of Condensin I (*elegans* Condensin Independent Domain or elCID). We investigate the progression of heterochromatin formation during development which reveals the early establishment of TADL domains and later appearance of compartments and elCID. These data reveal a diversity of structures that are established during early embryogenesis within nascent heterochromatin.

## Results

### Single-molecule Tracing defines well-insulated chromosome domains

To study heterochromatin organization, we focused on a chromosomal region characterized by a significant enrichment of methylated histone H3 at lysine 9 (H3K9me) ^39^, a well-known marker of the heterochromatic B compartment (Fig. 1A, Fig. S1A). This region was previously identified as part of the B compartment ^36^. HiC studies on *C. elegans* embryonic chromatin organization previously identified weakly insulated domains in this region, with poorly defined boundaries^23^. These domains had been defined using insulation scores for average matrices, which quantified the aggregate of chromatin interactions within a given genomic interval. Minima in the insulation profile, representing regions of high insulation were classified as TAD boundaries (Fig. 1A, Fig. S1A) ^23^. However, the lack of clearly visible domains with defined boundaries in averaged contact matrices raised questions about whether large-scale, canonical TADs existed in *C. elegans*^40^.

**Figure 1.**
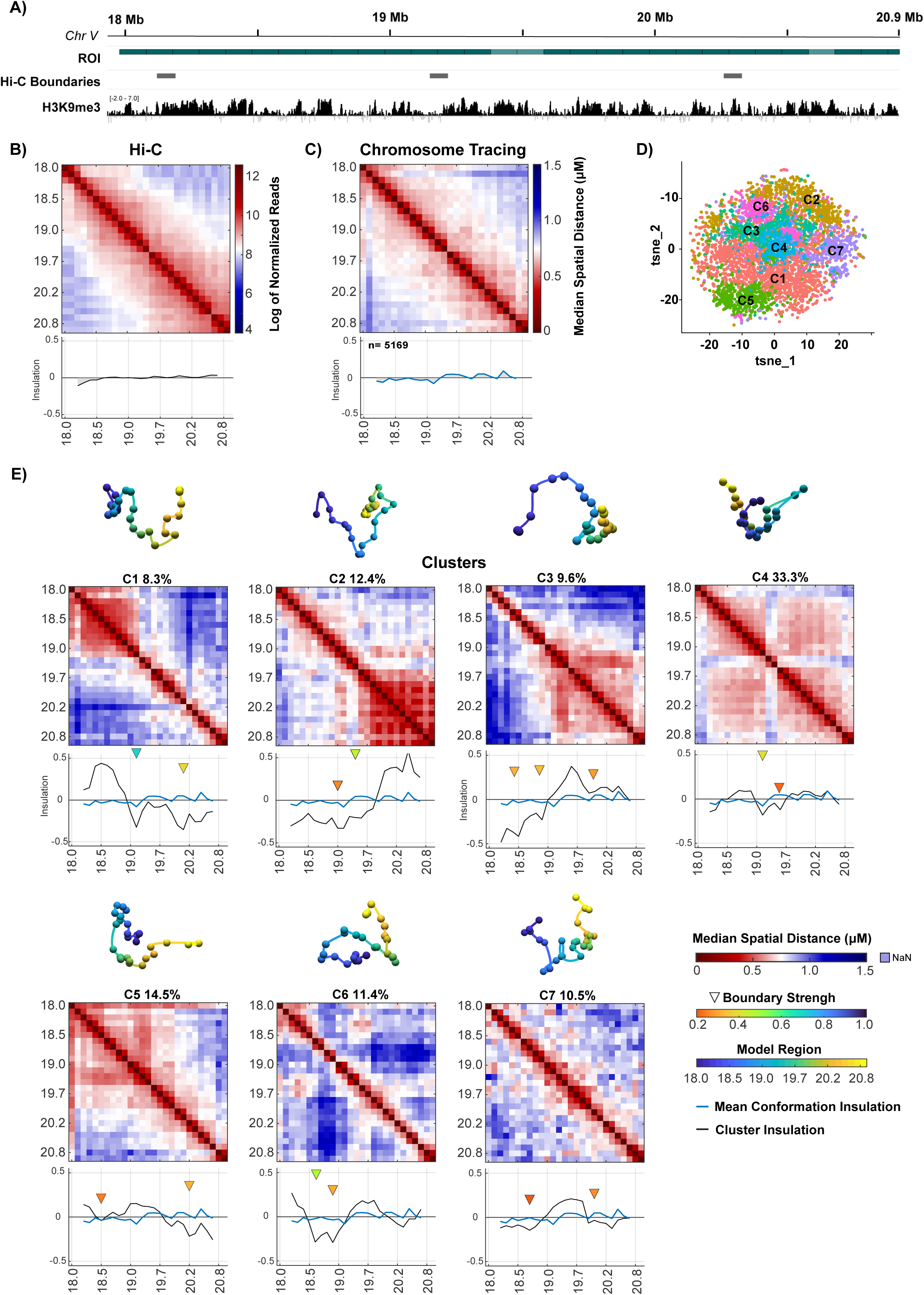
Single-molecule tracing reveals well-insulated chromosome domains in *c. elegans* embryos. **A)** Schematic of the Chromosome V region of interest (ROI) traced in this study (dark green); untraced flanking regions are shown in light green. Gray ticks mark Hi-C–defined TAD boundaries ^23^, and the H3K9me3 ChIP–seq signal is plotted below^39^. **B)** Log of normalized Hi-C sequencing read counts at 100 Kb resolution across the ROI with insulation score^32^. **C)** Median pairwise distance matrix (μm) for chromosomes at developmental stages spanning 2–140 cells with insulation scores. **D)** t-SNE plot of chrV clusters, where each dot represents a single chromosome trace and each color denotes a cluster. **E)** Median pairwise distance matrices (μm) for each cluster identified in (D), with insulation scores shown below. Triangles mark TAD boundary positions, colored by boundary strength. Above each matrix, three-dimensional bead-on-a-string models reconstructed by classical multidimensional scaling (MDS) illustrate cluster-averaged conformations; each sphere represents a genomic region positioned according to its median pairwise distances within the cluster.

We considered that domains might appear poorly defined due to population averaging of mixed populations, which included embryos at different phases of the cell cycle, distinct cell types and developmental stages. To address this issue, we employed Chromosome Tracing, a DNA FISH-based technique that enables direct observation of chromosome conformations at the single-molecule level ^36,37,41^. We focused on a heterochromatic H3K9me-rich region of Chromosome V^39^. In interphase nuclei, population-averaged Tracing data closely resembled Hi-C maps from mixed-stage embryos (r = 0.90; Fig. S1C), both lacking clear boundaries or strong domain structures (Fig. 1B–C), consistent with previous reports ^23,24,32,33^. These results indicate that the average chromatin conformation observed through Chromosome Tracing resembled that of Hi-C data, and both were characterized by weak domains and the absence of distinct boundaries.

To overcome the effects of population averaging and identify predominant chromosome configurations, we performed unsupervised clustering of single-molecule traces (Methods) ^36,41^. Additionally, we performed insulation analysis and calculated boundary strength at the single-molecule level, following the sliding box method of Crane et al. (2015). This analysis identified seven distinct conformational clusters (Fig. 1D–E), three of which exhibited ∼1 Mb domains with sharp boundaries and strong insulation scores (ranging from 0.5 to –0.4), nearly four times higher than the population average (Fig. 1E, C1–C3; Fig. S1D). We term these megabase domains. Clusters C1 and C2 showed the strongest boundaries and together accounted for ∼20% of all traces. We did not observe domains from Clusters C1 and C2 coexisting as separate structures within the same trace; rather, they appeared in an either/or configuration, or interacted to form larger domains resembling Cluster C4. The remaining clusters displayed weaker but above-average insulation values and distinct boundary positions, underscoring structural variability. These results demonstrate that *C. elegans* chromosomes adopt variable conformations at the megabase-scale, including a subset with domains with sharp boundaries. This structural heterogeneity and variable boundary positioning likely underlies the absence of clear domains in population-averaged data and aligns with the notion of a dynamically organized chromosome, consistent with recent findings from live imaging studies ^22,42,43^.

### Condensin is required for a subset of *C. elegans* heterochromatin domains

Genomic domains can be broadly categorized into two classes: those that rely on loop extrusion, such as TADs, and those that depend on attractive chromatin interactions, such as compartments ^2,8,19,21^. To determine the nature of our heterochromatin domains, we examined their genetic requirements. First, we examined the role of loop extrusion. While Cohesin in *C. elegans* forms small loops (∼20–40 kb) ^34,35^, Condensin I, an SMC complex, folds the genome at a larger scale ^23,24^. Given the megabase size of our domains, we focused on *dpy-28*, which encodes a subunit of Condensin I. Mothers that are mutant for *dpy-28(s939)* produce embryos that lack detectable DPY-28 protein, indicating a strong loss-of-function phenotype ^44^. Condensin II remains intact in *dpy-28* mutants, and likely explains the ability of *dpy-28* mutants to undergo mitosis and survive until larval stages^45^.

We began by examining the impact of Condensin I depletion at the population level (Fig. S2A). Median distance matrices from three independent replicates revealed that Condensin-deficient embryos displayed an alternating, plaid-like pattern instead of continuous domain-like interactions (Fig 2A-B, Fig. 2SB). In addition, nearly all pairs of probes in the region displayed significant larger distances, indicating the chromosomes were less compact (Fig. 2B, Fig. S2C). We observed increased proximity in two regions of the mutant traces, resulting in a reversal of the insulation profile (blue arrows Fig. 2C). This result indicates a gain of proximity and a corresponding loss of insulation upon DPY-28 depletion. Conversely, a region exhibiting low insulation in the wild type, (green arrow, Fig. 2C) became more insulated following Condensin I loss. RoG measurements confirmed a significant increase in chromosomal volume in *dpy-28* mutants compared with the wildtype (p = 1.05e-05; Fig. 2E), consistent with global decompaction (Fig. 2E).

**Figure 2.**
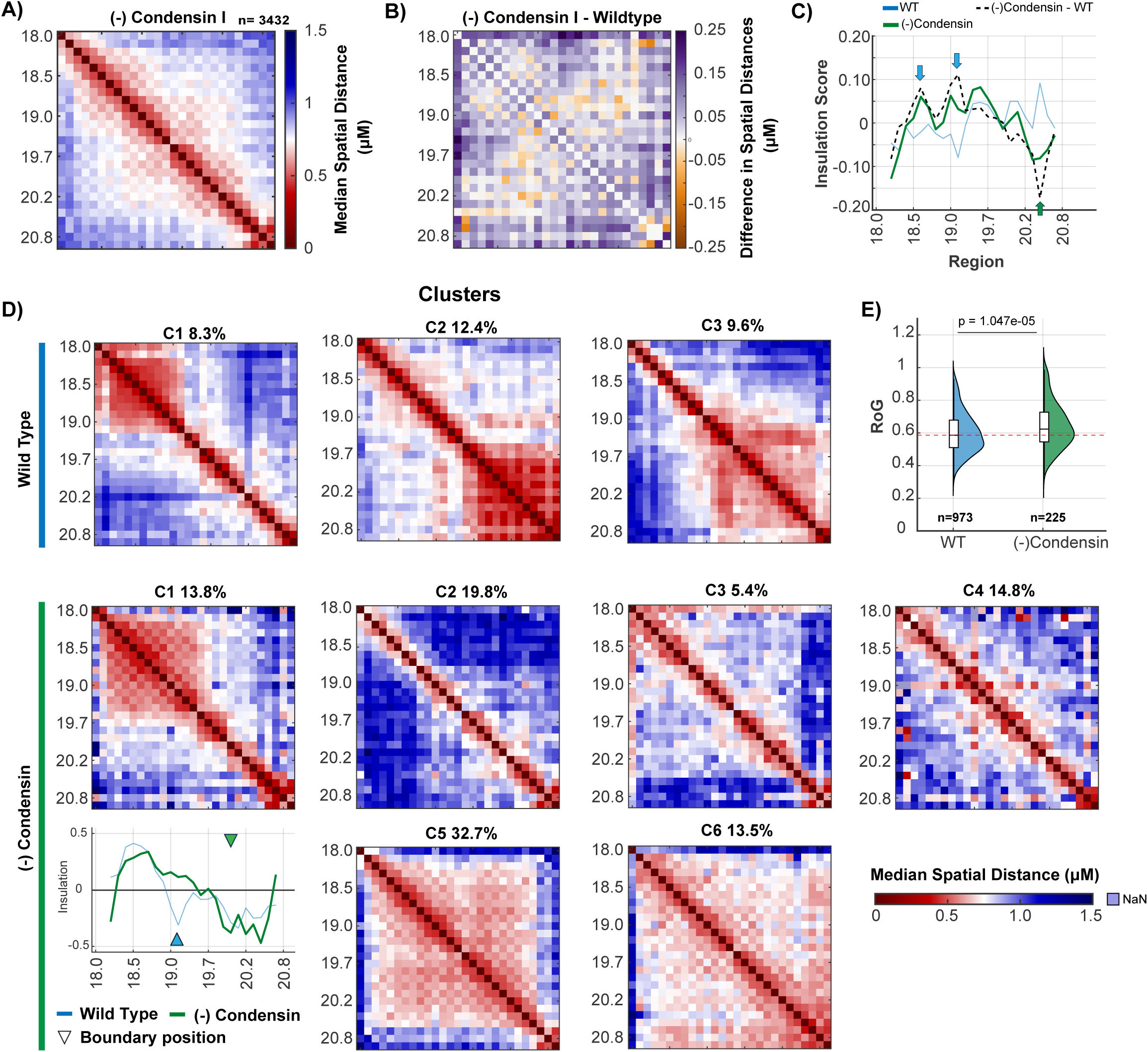
Condensin I is required for a subset of *C. elegans* heterochromatin domains. **A)** Median pairwise distance matrix (μm) for Condensin mutant from 2 to 140 cells, annotated with insulation scores. **B)** Difference in pairwise distances (μm) between Condensin(-) mutants and wild-type traces. Purple regions indicate increased distances in Condensin mutants, while orange regions indicate increased distances in wild-type. **C)** Insulation scores for wild-type, Condensin(-) average matrices and their difference. Blue arrows highlight regions with reduced insulation after Condensin depletion, while green arrows indicate regions with increased insulation. **D)** Median pairwise distance matrices (μm) for each cluster. The top row displays clusters with distinct domain boundaries in wild-type, while the bottom row shows clusters in Condensin(-) mutants. The insulation scores display the insulation of both C1 clusters in wild-type and Condensin(-) embryos. Green arrows mark boundaries in Condensin mutants; blue arrows mark boundaries in wild-type. **E)** Radius of gyration for wild-type and Condensin(-) traces. Statistical significance was determined using a t-test. n indicates the number of traces.

To understand the nature of our chromosomal region beyond the population level, we performed clustering using the same parameters as for the wild-type sample (Fig. 2D, Fig. S2D). We highlight clusters with the strongest boundaries in the wild type (Fig. 2D, *top row*). Our results showed that the two strong domains observed in WT-C1 and WT-C2 clusters were differentially affected by *dpy-28* mutations The distal domain spanning 19.7–20.9 Mb (C2) was lost, consistent with a requirement for Condensin I, whereas the proximal domain spanning 18.2–19.6 Mb (C1) persisted, though in a slightly decompacted form with weaker boundaries. We name this cluster elCID (*elegans Condensin-Independent Domain*), and note that this cluster contains more traces in Condensin mutants compared to the wild type (∼14% vs. ∼8%). Other mutant traces resolved into either highly open conformations (C2–C4; 40%) or compact states spanning the entire region (C5–C6; 46.2%) (Fig. 2D, bottom rows). Together, these results show that while some megabase domains, such as the distal TADL, strictly depend on Condensin I, others, like elCID, form independently, despite their resemblance to TADL.

### Histone H3K9 methylation mediates compaction but not boundary formation of heterochromatin domains

Epigenetic modifications contribute to compartmentalization but are not required for loop extrusion ^1,3,18,46–48^. To test whether C2 was a TAD-like structure, we hypothesized that its organization would remain largely unaffected by the removal of H3K9 methylation. To test this idea, we analyzed embryos lacking virtually all H3K9 methylation using *met-2; set-25* double mutants ^49,50^. Clustering of two independent replicates, using wild-type parameters, identified domain-containing clusters similar to those in wild type, including both elCID and the TADL domain (Fig. 3A, Fig. S3A, S3C). These findings indicate that megabase-scale domains still form in the absence of H3K9me. We conclude that these domains are not compartments, and that the domain at 19.7–20.9 Mb resembles a TADL domain with sharp boundaries, a dependency for an SMC factor and independence from H3K9me despite its location in a large region of heterochromatin.

**Figure 3.**
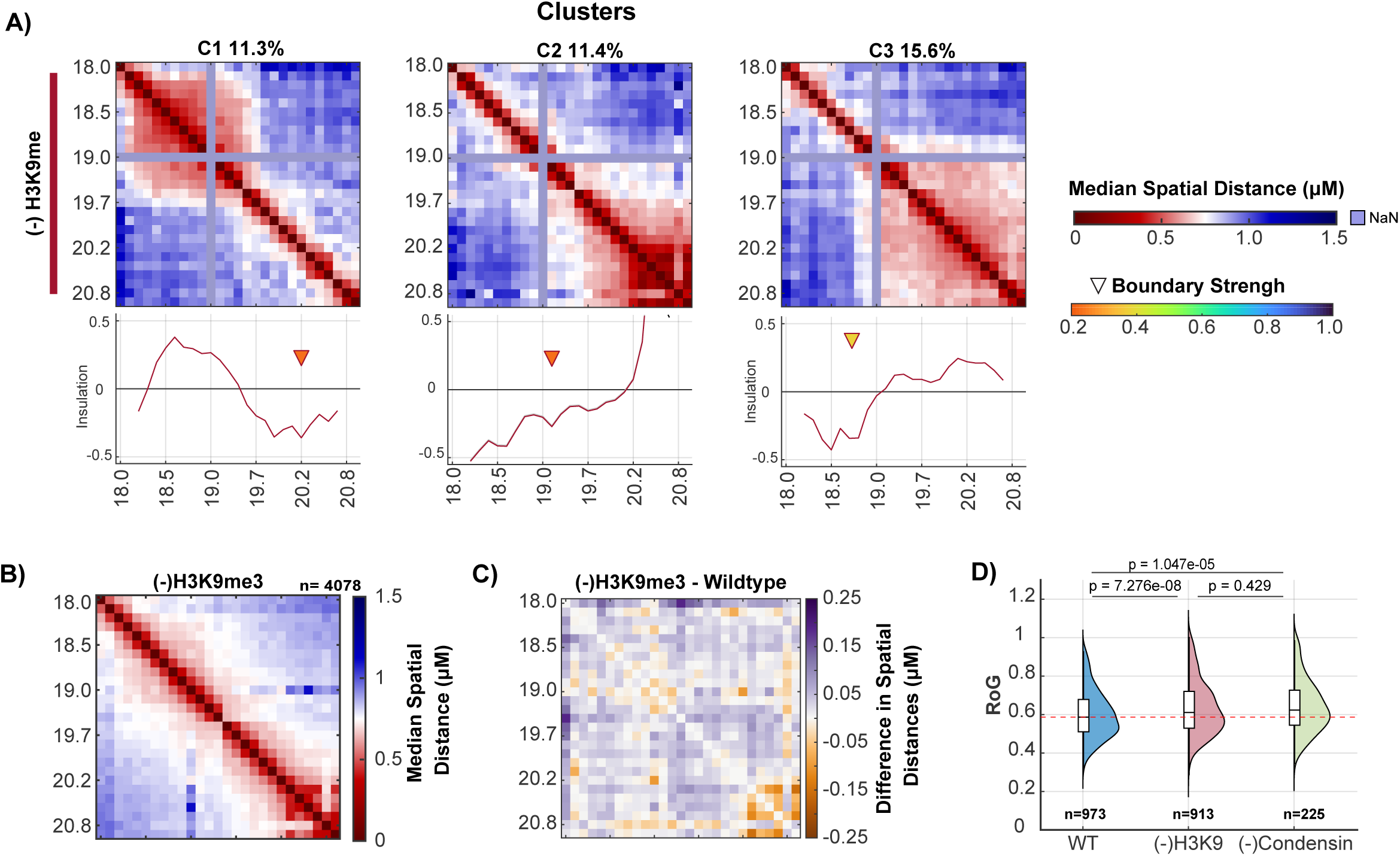
H3K9me mediates compaction but not boundary formation of heterochromatin domains. **A)** Median pairwise distance matrices (μm) for C1, C2, and C3 clusters in H3K9me traces, annotated with insulation scores. Triangles indicate TAD boundary positions, with colors representing boundary strength. **B)** Median pairwise distance matrix (μm) for H3K9me(-) mutant chromosomes across developmental stages from 2 to 140 cells **C)** Differences in pairwise distances between H3K9me(-) mutants and wild-type traces. Purple regions indicate distances larger in H3K9me(-) mutants, while orange regions indicate distances larger in wild type. **D)** Radius of gyration for wild-type, H3K9me(-), and Condensin(-) traces. Statistical significance was determined using a t-test. n indicates the number of traces.

H3K9 methylation is crucial for compacting chromosomes during the formation of large-scale heterochromatin ^46,50,51^. To evaluate its effect, we measured median distances and the radius of gyration (RoG) in wild-type and *met-2 set-25* mutant embryos. As expected, mutants showed significantly increased distances across the region, indicating reduced chromatin compaction (Fig. 3B–C, Fig. S3D). RoG analysis confirmed significantly larger chromosome volumes in mutants (p = 7.28e-08), comparable to those in *dpy-28* embryos (p = 0.429) (Fig. 3D). Down sampling controls ruled out bias due to sample size (Fig. S3E–F). These results show that loss of H3K9 methylation causes global chromatin decompaction, affecting both Condensin-dependent and independent domains.

### Distinct Genetic Dependencies Shape TAD-like and elCID Chromatin Structures

Clustering chromosome traces independently by genotype does not reveal whether similar clusters reflect equivalent structures across genotypes (e.g. C1 in wild-type vs C1 in H3K9me mutants). To identify genotype-specific structures, we pooled chromosome traces from wild-type and mutant samples, and clustered the mixed population (Fig. 4A). To ensure unbiased comparisons, we down-sampled the datasets to achieve uniform age distributions across the three genotypes (*Fig. S4A-B*, *see Methods*). We note that no differences were observed for single genotypes between the down-sampled and full datasets, confirming the robustness of the analysis (Fig. S4B).

**Figure 4.**
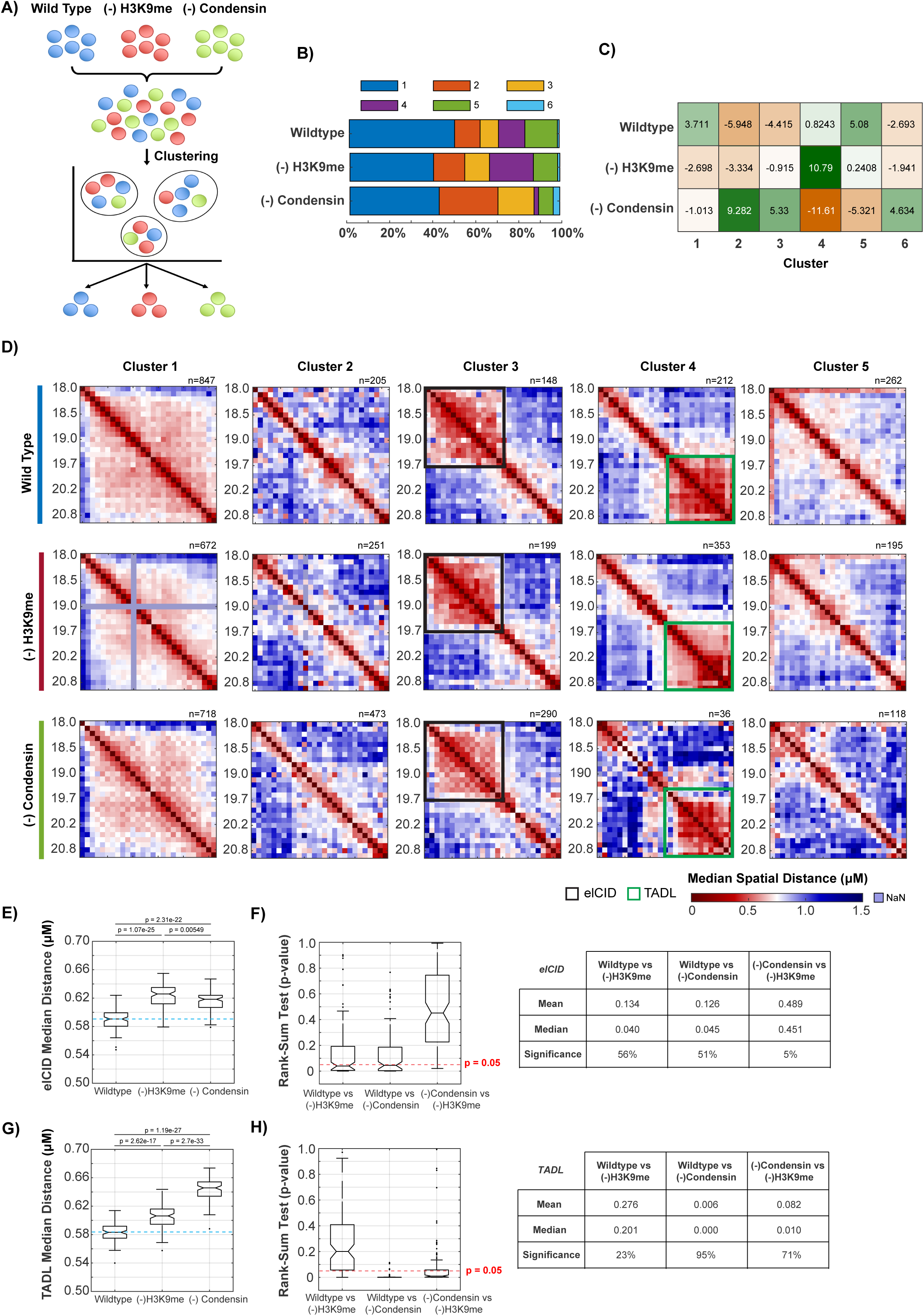
Chromosomal clustering highlights condensin’s essential role in tad-like organization. **A)** Schematic representation of the co-clustering strategy used to group chromosome traces based on structural similarities. **B)** Contribution of each genotype to the clusters identified by doing (A), illustrating how different genotypes are distributed across structural clusters. **C)** Standardized chi-square residuals, as described in Gutnik et al., 2024^41^. High positive values indicate enrichment of traces from a specific genotype in a given cluster, while negative values indicate depletion of traces in that cluster. **D)** Median pairwise distance matrices (μm) for each identified cluster. Co-clustering was performed by pooling chromosome traces from different genotypes. **E–H)** Iterative down-sampling analysis of elCID (E, F) and TADL (G, H) spatial distances across wild type, H3K9me(–), and Condensin(–) conditions. For each dataset, 1,000 distances were randomly sampled across 100 iterations. **E, G)** Boxplots of median spatial distances per condition; horizontal bars mark medians and p-values from full datasets are shown above. **F, H)** Corresponding boxplots of p-values from pairwise Wilcoxon rank-sum tests across the same iterations, with the red dashed line indicating p = 0.05. Adjacent tables report mean and median p-values and the fraction of iterations with p < 0.05, reflecting the robustness of differences observed in (E, G).

Clustering the mixed population revealed genotype-specific enrichment or depletion for certain clusters (Fig. 4B–D, Fig. S4C–F). The most pronounced difference was seen in Cluster C4, corresponding to the TADL domain. This cluster was present in the wild type, enriched in H3K9me mutants, but nearly absent in *dpy-28* mutants (chi-square residual = –11.61), indicating a strong dependence on Condensin. Cluster C3 corresponded to the domain at 18.2–19.6 that was enriched in Condensin mutants (chi-square residual = 5.3) and unaffected by H3K9me loss. We presume this is elCID, highlighting that domains with similar features can arise through distinct regulatory factors. Finally, Cluster C2, which showed a decompacted conformation, was significantly enriched in *dpy-28* mutants (chi-square residual = 9.3) and decreased to about half in wild type and H3K9me mutants.

Across all three genotypes, the predominant chromatin configuration was Cluster C1, a highly interactive domain spanning the entire region, with intermediate folding and no clear internal boundaries (Fig. 4D). This cluster accounted for ∼40% of traces per genotype, indicating that loosely organized structures are common. Subtle genotype-specific differences were evident: in *H3K9me* mutants, chromatin proximity declined more sharply with genomic distance, consistent with a role for H3K9me in stabilizing long-range interactions. In *dpy-28* mutants, we again observed the plaid-like pattern, which was also visible in compact domains such as C3. A similar but weaker plaid signature was detected in wild type, whereas *H3K9me* mutants exhibited a smoother version of this pattern. Together, these results reveal the genetic requirements of distinct domains at larger and also finer scales.

To compare elCID and TADL, we measured inter-region distances within their respective domains on chromosome V (elCID: 18.2–19.6 Mb; TADL: 19.7–20.8 Mb) in wild-type and mutant embryos (see Methods, Fig. 4E–H). The two domains showed contrasting behaviors. elCID displayed only modest increases in median distances (0.62 µm vs. 0.59 µm in wild type; Fig. 4E–F), and these effects were small and inconsistent across subsampling iterations, suggesting relative stability. In contrast, TADL showed pronounced and reproducible decompaction (Fig. 4G), with significance maintained across nearly all iterations (Fig. 4H). Together, these findings show that while TADL behaves like a canonical, SMC-dependent TAD, elCID represents a robust alternative folding mode that persists independently of Condensin.

### Heterochromatin domains and domain compaction emerge progressively during development

To examine how heterochromatin domains arise during development, we analyzed chromosome structure across five defined embryonic stages: 2–4 cells (pre–zygotic genome activation, ZGA), 5–8 cells (minor ZGA), 9–40 cells (gastrulation onset, first major ZGA), and 41–80 and 81–140 cells, coinciding with the progressive accumulation of H3K9 methylation ^50,51^. We used co-clustering and power-law fitting and scaling analysis to quantify chromatin compaction over time.

Before ZGA (2–4-cell stage), the TAD-like domain (TADL) was already detectable in ∼30% of wild-type and H3K9me(-) embryos, but nearly absent in *dpy-28* mutants, indicating this domain appeared very early and presumably independently of transcription (Fig. 5A, Fig. S5A-B). In contrast, the Condensin-independent elCID domain emerged later (9–40-cell stage) across all genotypes, then disappeared, suggesting transient formation and distinct regulation (Fig. S5C). This timing aligns with previous EM studies showing the emergence of heterochromatin at gastrulation ^50,51^supporting elCID as a potential precursor to higher-order heterochromatin compartments.

**Figure 5.**
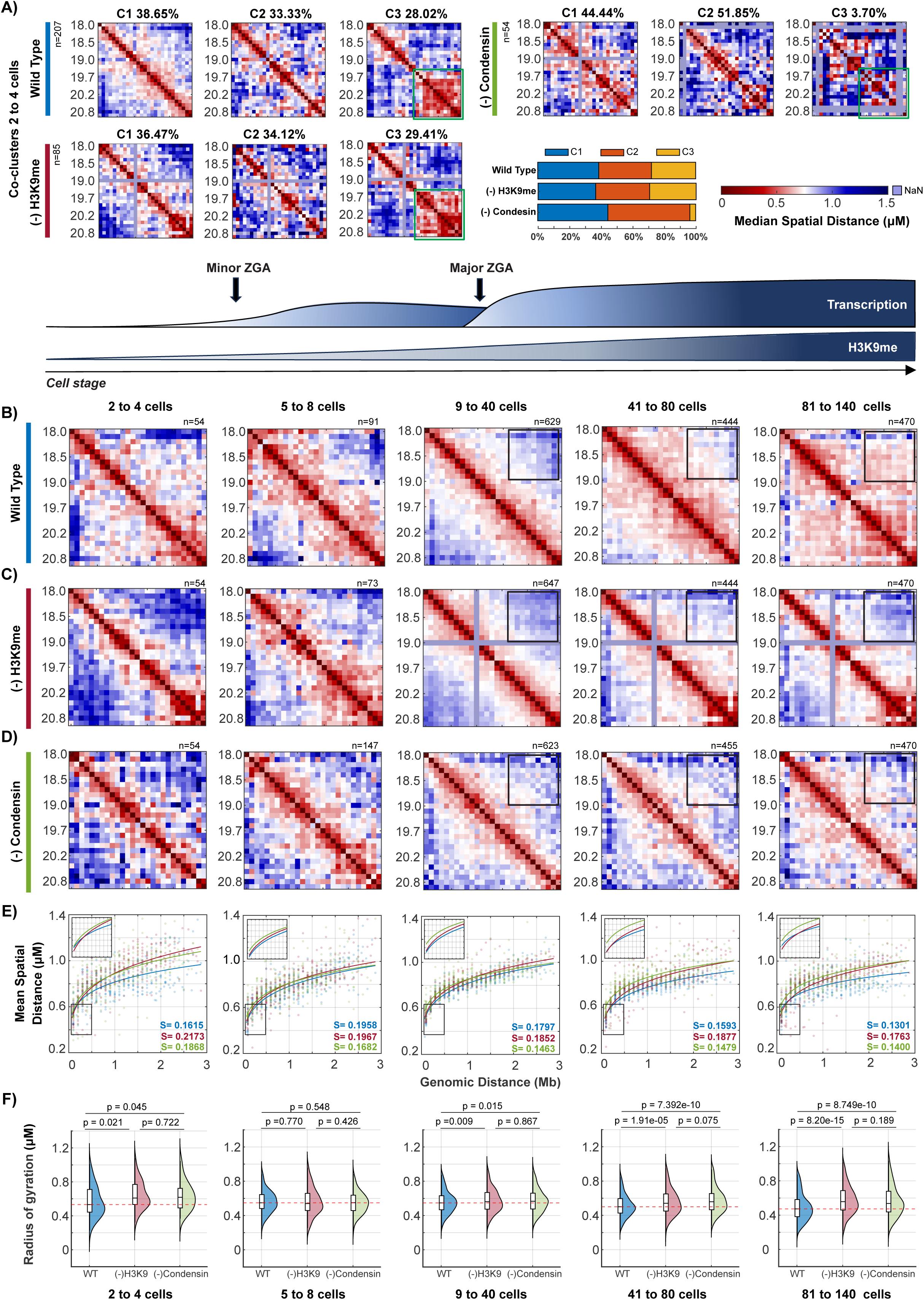
Onset of heterochromatin formation, chromatin compaction, and higher-order domain organization during development. **A)** Median pairwise distance matrices (µm) for embryonic co-clusters at the 2–4 cell stage, separated by genotype as in Fig. 4A. Left, contribution of each genotype to the clusters. **B–D)** Median pairwise distance matrices (µm) across five developmental stages for each genotype. *n* indicates the number of traces. Black boxes highlight long-range interaction regions that change during development. **E)** Scaling of mean pairwise distance versus genomic distance for wild-type (blue), H3K9me(-) (red), and Condensin(-) (green) across the five developmental stages. The black box provides a zoomed-in view of distances below 0.5 Mb (shown in Supplementary Figure S5A). **F)** Radius of gyration for wild-type (blue), H3K9me(-) (red), and Condensin(-) (green) across the five developmental stages. Statistical significance was determined using a t-test.

Chromatin compaction progresses during development in the wild type, but failed in both H3K9me(–) and Condensin(–) embryos. To quantify this folding, we analyzed the scaling of mean spatial distances with genomic distance, which follows a power law (*R* = *a*N^s^*), where the scaling exponent (s) reflects folding and the step size (a) reflects packing density. In wild type, scaling exponents declined from the 4-cell stage (s = 0.19 → 0.13; Fig. 5E, Fig. S6A-B), consistent with progressive compaction. H3K9me mutants remained elevated across development, indicating persistent decompaction. Condensin mutants, by contrast, showed lower exponents than wild type early in development (s = 0.16 vs. 0.19) but converged to wild-type levels by 80 cell stage. The step sizes remained higher in both mutants at 1 Mb, reflecting expanded chromatin, and only Condensin mutants showed elevated step sizes at 100 kb, pointing to a unique defect in local folding (Fig. S6A-D). Radius of gyration measurements confirmed these defects, with both mutants showing enlarged chromosome volumes from the 9-cell stage onward (Fig. 5F). Together, these results show that in the wild type, elCID and the TADL domain are compacted and may form a joint compartment over time (Fig. 5B).

Lastly, we tested whether these chromatin defects could be explained by changes in nuclear size. Across development, H3K9me(-)embryos maintained nuclear volumes comparable to wild type, suggesting that the observed chromatin changes are not due to differences in nuclear scaling (Fig. S6E). In contrast, *dpy-28* embryos exhibited significantly smaller nuclei, despite increased chromatin volume. This unexpected result indicates that the effects we see are not secondary effects due to nuclear size.

### Single-Cell Chromatin Domains Persist Despite Loss of Population TADs and Arise from Cooperative Long-Range, Not Free Polymer, Interactions

A puzzling attribute of TADs in other animals is that TAD-like structures can be observed in single-molecule traces of SMC mutants but vanish in population-averaged maps^47,52^. We examined single-molecule traces in wild-type and mutant embryos, and observed a similar phenomenon (Fig. 6A,C). We measured the boundary probability, defined as the likelihood that two neighboring bins are separated by more than a threshold distance (Methods). In the wild type, two regions stood out with elevated boundary frequencies, at 19.0 Mb (∼20%) and 19.7–19.9 Mb (∼15%) (Fig. 6B), consistent with the locations separating the elCID and TADL domains. We note that these boundary probability percentages are consistent with values reported in human cells ^47,52^. We also observed a range of structures, by visual inspection, revealing the diversity of possible folds (Fig. 6a). In *dpy-28* mutants, domain-like structures remained detectable in single Condensin(-) traces (Fig. 6C, Fig. S7B). However, boundary probabilities were evenly distributed across the region (Fig. 6D), indicating increased variability in boundary positioning across cells. In H3K9me mutant embryos, boundary positions remained centered near 19.0 and 19.8 Mb, similar to wild-type, with slightly reduced probabilities (Fig. S8A–B). elCID domains in these embryos showed increased interactions near boundary regions (Fig 4D, C3), but overall boundary strength was not significantly changed (Kolmogorov–Smirnov test, p = 0.89; Fig. S8C), suggesting a minor role for H3K9me in reinforcing boundaries. These results show that chromatin domains persist at the single-cell level in the absence of Condensin or H3K9me, similar to other animals ^47,52,53^.

**Figure 6.**
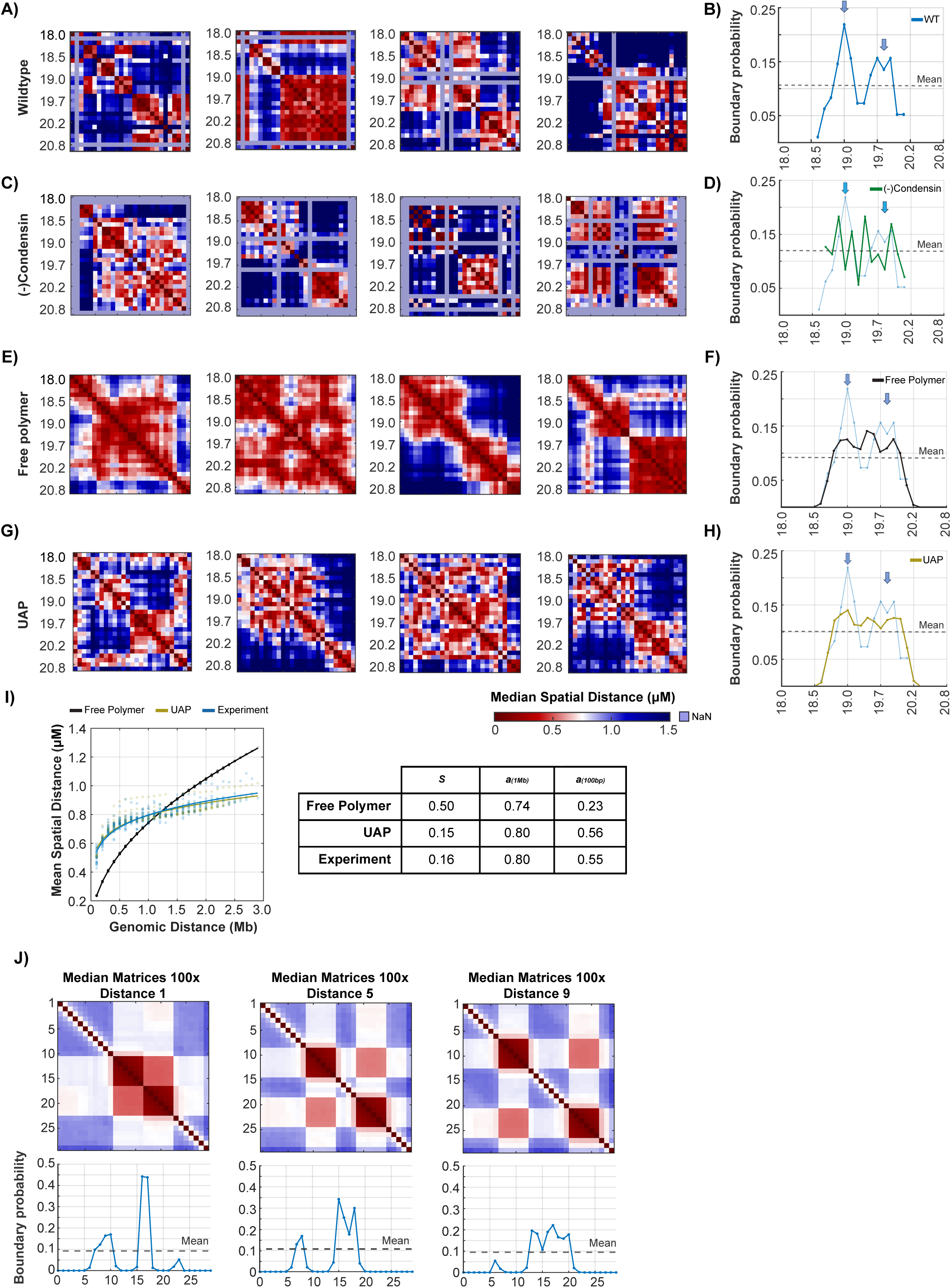
Chromatin domains require cooperative and long-range interactions beyond polymer physics. **A, C, E, G)** Representative examples of single-chromosome spatial distance maps (µm) for wild-type embryos (A), Condensin-depleted embryos (C), free polymer simulations (E), and uniformly attractive polymer (UAP) simulations (G). **B, D, F, H)** Boundary probability profiles derived from collections of single-chromosome traces for the corresponding conditions. Boundary probability reflects the fraction of single traces in which a genomic bin is classified as a boundary (see Methods). Arrows in boundary plots highlight regions with high boundary probability in wild-type. Dashed lines indicate the mean boundary probability across the region. **I)** Scaling relationships between genomic and spatial distance for experimental data, free polymer, and UAP simulations. The table shows the estimated scaling exponent (*s*) and step sizes (*a*) at 1 Mb and 100 kb for each condition. **J)** Median distance matrices from simulations of two size-6 domains with strong intra-domain interactions (100×), separated by 1, 5, or 9 regions Boundary probability profiles (bottom) show that closer spacing reinforces boundary strength, while wider spacing produces weaker, broader peaks.

We considered that single molecule domains might reflect intrinsic polymer fluctuations, which we modeled with a minimal polymer model by modeling polymers as a Gaussian process ^54,55^. In this framework, an interaction matrix defines pairwise contact strengths, and ensembles of configurations are drawn from a multi-variate normal distribution, capturing both structured and stochastic folding behavior. To evaluate how much chromatin folding could be explained by minimal polymer interactions, we simulated two distinct models, each generating 10,000 conformations (Fig. S8D). In the simplest case, we used a free polymer model, which included only nearest-neighbor interactions and was calibrated to match the experimental radius of gyration (Fig. S8E–F). At the single molecule level, the free polymer model produced molecules with strong insulation and boundaries (Fig. 6E). This result suggests that some structures seen in vivo, particularly in Condensin mutants, could reflect free polymer behavior. However, the computed biophysical properties of these structures were dramatically different from those measured in vivo (Fig. 6I). The scaling exponent (s) in wild-type embryos was markedly lower than that of a free polymer (s = 0.16 vs. 0.50), indicating long-distance interactions and significantly greater overall compaction in vivo.

To incorporate long-range, distance-dependent interactions, we implemented a Uniformly Attractive Polymer (UAP) model (Fig. S8D–G). UAP simulations produced more compact and entangled structures, with heterogeneous, fragmented domain-like features resembling those observed in single-cell data, in contrast to the smooth domains of the free polymer (Fig. 6A, D; Fig. S7A, D). This model more accurately reproduced wild-type biophysical parameters, including a shallow scaling exponent (s = 0.15) and step sizes of a = 0.56 µm at 100 kb and a = 0.90 µm at 1 Mb (Fig. 6I). However, while the UAP model recapitulated overall compaction, it failed to reproduce boundary probabilities (Fig. 6H), underscoring that unstructured interactions that are homogeneous along the polymer, as assumed in the UAP model, are insufficient to account for domain architecture.

We constructed models with heterogeneous nearest-neighbor interactions (enhanced 3×, 10×, or 100×; Fig. S8H) to test how local contacts generate domains. Stronger interactions compacted domains in population averages (Fig. S8I), but only very strong interactions (100×) produced robust single-molecule domains and appreciable boundary signals (Fig S8J,L), indicating that intra-domain contacts must reach a high threshold to generate both domain structure and stable boundaries. We next examined how adjacent domains influence boundary formation by simulating chromosomes with two size-6 domains stabilized by strong intra-domain interactions (100×) and varying their spacing (1, 5, or 9 regions; Fig. S8K). In population-averaged matrices, the two domains frequently associated despite lacking explicit inter-domain attraction (Fig. 6J), though this association was variable at the single-molecule level (Fig. S8M). Boundary probability profiles revealed that close domains reinforced one another: when separated by a single region, boundary signals merged into a sharp peak reaching 44%, whereas wider separations produced only a weaker, broad plateau (Fig. 6J). Thus, while strong intra-domain interactions can generate compact domains, stable boundaries require reinforcement between neighboring domains, explaining why boundary probability is lost in condensin mutants when the TADL is absent even though elCID persists.

To further assess how well minimal polymer models capture the structural diversity of in vivo chromatin, we co-clustered simulated and experimental single-chromosome traces (Fig. S9). The free polymer model primarily produced compact but unstructured configurations, corresponding to only a subset (∼25%) of in vivo traces (Fig. S9A-C). The remaining clusters were largely model or wildtype-specific, indicating structural differences not captured by polymer physics alone. In contrast, the UAP model better recapitulated experimental variability: 73.5% of simulated traces clustered with C1, composed of compact structures, while 19.2% aligned with domain-containing clusters (C3 and C6; Fig. S9B-D). However, the TAD-like cluster (C4), which accounted for 15% of experimental traces, was underrepresented in UAP simulations (2.5%), suggesting that uniform long-range attraction alone cannot generate Condensin-dependent domains. These results demonstrate that while distance-dependent interactions reproduce general folding behaviors, additional mechanisms, such as Condensin-mediated boundary formation are required to fully account for the diversity of single-cell chromatin conformations.

## Discussion

The diversity of chromatin domains that package chromosomes remains a central question in genome biology. Our study addresses this issue by identifying megabase-scale domains in *C. elegans* heterochromatin and dissecting their genetic and biophysical features. We identified TADL-like domains that were highly folded, display boundary probabilities comparable to those in mammalian cells ^47,52,53^, and required Condensin I for their formation. We also identified elCID, a domain that resembled the TADL but was largely independent of Condensin I. Though dependent on different factors, these domains converge to promote compartment-like heterochromatin.

We observed TAD-like domains prior to ZGA, indicating that these domains initiate independently of transcription. Earlier studies in other organisms have reported conflicting results regarding the relationship between TADs and ZGA, with some observing a need for transcription and others not ^56–60^. Our findings suggest that these discrepancies may arise from population-averaged approaches, which can mask structures that exist for only a subset of chromosomes. In *C. elegans*, the TADL cluster represented a substantial fraction of traces prior to ZGA (∼28%), but still a minority.

Previous studies suggested that *C. elegans* lacks TADs because this species lacks CTCF and domains with strong boundaries ^23,24^. However, our study suggests this conclusion reflects variability rather than absence of defined domains, despite the lack of an obvious CTCF orthologue ^61^. We observe TAD-like domains that exist at the single-molecule level, are resolved by clustering, and require Condensin I. In other systems, boundary formation often involves multiple cooperating insulator proteins, many of which are zinc finger–containing factors^62–68^. The *C. elegans* genome encodes numerous zinc finger proteins, raising the possibility that some act as alternative insulators.

Beyond the elCID and TADL domains, we also observed less defined structures, and these made up the majority of traces. Recent studies with mammalian cells indicate that TADs reflect only a small proportion of chromosome structures in vivo (≤3%)^22,42^, suggesting that vertebrate chromosomes could likewise be dominated by less structured conformations. Such configurations are difficult to capture by Hi-C, which requires chromatin segments to lie within <10 Å for crosslinking, but become evident through single-molecule, imaging approaches. Future studies with vertebrates will determine whether comparable structural diversity exists in other species.

Within the megabase domains, we detected intermittent, plaid-like patterns in Condensin mutants, and to a lesser degree in wild-type embryos. The absence of this pattern in H3K9me mutants suggests that they are not artifacts of the tracing pipeline, but instead represent biologically relevant structures. Such plaid pattern could reflect increased chromatin flexibility and abnormal interactions following SMC loss ^69^. This possibility could also explain the lack of plaid structures in H3K9me mutants, where compartmental interactions are weakened^33^. Alternatively, residual or aborted Condensin activity, especially *in dpy-28* mutants, might generate smaller loops that appear plaid-like.

## Materials and Methods

### Worm strains

*C. elegans* strains used in this study are: N2 Bristol strain (wild-type), GW638 *met-2(n4256) set-25(n5021)* III and TY4381 *dpy-28(s939)/qC1 [dpy-19(e1259) glp-1(q339) qIs26]* III. All strains were cultured at 20 °C on NGM plates seeded with OP50 grown in LB.

### Genomic data

#### Hi-C data

Hi-C data for *C. elegans* embryos, accession number GSE206065 ^32^ was used to generate contact matrices at a 100 Kb resolution. The data was normalized using the ICE (Iterative Correction and Eigenvector decomposition) method ^70^, with regions excluded from tracing similarly omitted from the Hi-C plots. To evaluate the correspondence between Hi-C and tracing data, a Pearson correlation analysis was performed, revealing a strong correlation between the averaged spatial distances and the normalized Hi-C contact frequencies (Supplementary Fig. 1a). Hi-C boundary information for Fig. 1A was downloaded and plotted from Crane et al. (2020)^23^, with the accession number GSM1556154.

#### CHIP-seq data

ChIP-seq data for H3K9me3 (accession number GSE22720^39^) was downloaded and analyzed using IGV for visualization and plotting.

#### Primary Probe design

Probe design and synthesis methods were as previously described by ^38^. DNA sequence was extracted from *C. elegans* genome assembly (ce11). Primary probes, each 102 nucleotides in length, target consecutive 100 kb regions covering the distal right arm of chromosome V. Each primary probe sequence contains 4 parts: 5’ PCR-priming sequence (20 nt), Genome homologous sequence (42 nt), readout sequence (20 nt), and 3’ PCR-priming sequence (20 nt) from ^71^. The OligoArray 2.1 ^72^ parameters used were: targeting region length 42 nt, melting temperature 78-100°C, no internal secondary structure with melting temperature greater than 76°C, no cross-hybridization with melting temperature greater than 76°C. GC content 30%–90%, and sequences do not contain consecutive repeats of seven or more identical nucleotides.

OligoArray2.1 was run on sciCORE for high-performance computing at University of Basel. The 42 nt homologous sequences were further blasted by NCBI BLAST+ 2.9.0 for specificity over the *C. elegans* genome.

### Chromosome tracing

Tracing was conducted with probe sets at 100 Kb resolution, using Chromosome Tracing as described in Sawh et al. (2020)^36^ and Wang et al., (2016)^37^. Below is a general description of the protocol employed to conduct the experiments.

#### Sample preparation

Gravid young adult worms were dissected in ddH₂O to isolate embryos. The embryos were then transferred onto round cover slips pre-coated with Poly-L-Lysine for better adhesion. Embryos were treated with a solution of 1% paraformaldehyde and 0.05% Triton X-100 for 5 minutes to fix them. The slides were placed on dry ice and freeze-cracked ^38^.

For *dpy-28(s939)*, homozygous adults were selected based on their lack of GFP. Mutant embryos were collected from homozygous mutant mothers, ensuring they were free from maternal contribution (Supplementary Fig. 3a) ^44^.

#### Hybridization Protocol

After freeze-crack fixation, samples were rinsed with phosphate-buffered saline (PBS) and washed three times with 1X PBS containing 0.5% Triton X-100. Following these washes, embryos were incubated with RNase A (0.05 mg/ml) for 30 minutes at 37°C to digest RNA. After the incubation, the RNase was removed, and the samples were placed in a hybridization solution composed of 2X SSC (Saline-Sodium Citrate buffer) with 10% dextran sulfate, 0.1% Tween-20, and 50% formamide, and incubated at 47°C for 1 hour.

Next, primary probes were diluted in hybridization buffer at 1:100, added to the samples and incubated for 10 minutes, before transferring to a hot metal surface at 85°C for 10 minutes. Next, the slides were moved to a humid chamber for probe hybridization, which was allowed to proceed for at least 16 hours at 47°C. Post-hybridization, the slides were washed for 1 hour with 2X SSC in 50% formamide, followed by two washes in 2X SSC and two in 0.5X SSC at 47°C, each lasting 30 minutes. The slides were either imaged immediately or stored at 4°C.

For secondary hybridization, complementary probes were diluted to a concentration of 8 nM in a solution consisting of 2X SSC and 25% ethylene carbonate (Sigma Aldrich). The samples were incubated with these secondary probes for 30 minutes, followed by a wash for 3 minutes in the same 2X SSC and 25% ethylene carbonate solution at room temperature, and a subsequent wash for 3 minutes in 2X SSC alone at room temperature Images of the secondary probes were captured using 561 nm or 647 nm channels. In the sequential rounds of hybridization, oligonucleotides were introduced to displace the previously bound secondary probes and were incubated for 30 minutes alongside the secondary probes targeting the subsequent region.

#### Microscope imaging pipeline

Experiments were performed using a widefield microscope (Ti2-Crest NIKON) integrated with a custom microfluidics system at room temperature, where secondary probes were delivered to the sample ^36,41^. Prior to imaging, embryos were stained with DAPI, and fiducial beads (488 nm) were added to the sample. Fields of view (FOVs) were selected for imaging, during which both the primary probe and DAPI signals were captured. Following this, the primary probe was subjected to photobleaching. Sequential imaging rounds for the secondary probes were conducted after hybridization. Tetraspeck fluorescent beads were imaged in every imaging round to enable accurate channel alignment

#### Imaging analysis

After tracing, the acquired images underwent foci fitting as previously described ^36,37^. For image segmentation and chromosome tracing, MATLAB was utilized, implementing modified methods outlined in prior studies ^36,41^. Initially, DAPI fluorescence was employed to identify nuclei within each embryo. Background signals were subtracted, and noise was removed. The images were then converted to binary format, distance transformed, and processed using watershed segmentation, resulting in volumetric masks that delineated individual nuclei. Chromosome territories were segmented in a similar fashion, with the nuclear mask excluding signals outside the nuclei. This process produced a volumetric representation in which each volume corresponded to a distinct chromosome. Chromosome tracing was subsequently executed within these defined volumes using the nearest-neighbor approach as previously described by Sawh et al. (2020). Finally, we reconstructed median distance matrices from the chromosome tracing data, using a minimum of seven detected regions averaged across replicates, which demonstrated a strong correlation. Regions 19.4, 19.5, and 20.6 were excluded from the analysis due to the absence of detectable hybridization signals. Consequently, these regions are not shown in the plots. In the H3K9me(-) datasets, region 19.0 contains an artifact in the matrix caused by low hybridization efficiency, leading to a reduced number of measurements in this region.

### Chromosome conformation Analysis

#### Pearson correlation between replicates

For each genotype, a Pearson correlation coefficient was calculated to assess the mean pairwise spatial distance consistency between regions and across replicates. The MATLAB function *corrplot* was employed for these analyses.

#### Visualization of Individual Traces

To visualize individual genomic region traces in three dimensions, we utilized the scatter3 function in MATLAB. Each trace was represented in a 3D space, with the X, Y, and Z parameters corresponding to the coordinates of each point in the trace. Input vectors for these coordinates were provided to the function, allowing for the representation of spatial relationships among the regions. The points were connected with lines to illustrate the trace path, facilitating the examination of the dynamic interactions among the genomic regions.

#### Insulation Analysis

We calculated insulation scores using the method described by Crane et al. (2015)^23^, with modifications for our chromosome tracing data. First, we normalized the mean distances to account for genomic distances, particularly considering the presence of non-consecutive hybridization regions, which could introduce potential biases in the data. To achieve this normalization, we fitted the data to a power-law function to accurately reflect the expected relationship between spatial proximity and genomic distance ^37^. We employed a sliding window of 200 kb along the diagonal of the interaction matrix to average the number of interactions across these bins. The mean signal within each window was then assigned to the corresponding chromosome tracing region. Bins located at the beginning and end of the matrix were excluded from the analysis, as they extended beyond the matrix boundaries. It is important to note that no data were available for the first and last regions due to the insulation box constraints.

We normalized insulation scores for each chromosome by computing the log2 ratio of each bin’s score to the mean insulation score across all bins of that chromosome. This normalization allowed comparison of local insulation relative to the overall chromatin organization within the chromosome. Local minima (“valleys”) in the normalized insulation profile were taken to represent regions of high insulation. To systematically identify these valleys, we calculated a delta vector, defined as the difference between the mean insulation values within 100 kb upstream and 100 kb downstream of the central bin. Valleys were detected at zero-crossings of this delta vector. For each boundary bin, we quantified boundary strength as the difference in delta values between the local maximum to the left and the local minimum to the right, following the approach of Crane et al. (2015) ^23^. Bins with boundary strength ≥ 0.1 were classified as boundaries.

#### Boundary probability

To calculate the boundary probability of the traces, we selected those with a minimum of 25 regions and generated a mean distance matrix. Linear interpolation was applied between maximum consecutive 2 regions to address any missing data points, and the data was then normalized using the average values obtained. Insulation analysis was conducted on individual traces, following the previously described method, with an insulation box of 300 kb and a delta box of 200 kb. Bins with boundary strength ≥ 0.5 were classified as boundaries.

After obtaining the boundaries for all the traces, we calculated the probability across all regions, similar to the approaches outlined by Cheng et al. (2021) and Bintu et al. (2018)^47,52^. To compare the distribution of boundary strength and probability between genotypes, we employed the Kolmogorov-Smirnov Test (Two-Sample K-S Test). This statistical test evaluates the cumulative distributions of two datasets to determine whether they have same distribution.

#### Clustering

Clustering was conducted as described in Sawh et al. (2020)^36,41^. Spatial distance matrices were constructed for individual chromosome traces by vectorizing the unique pairwise measurements. These distance vectors were then concatenated into a single file, linking each trace to its original file containing the foci xyz coordinates. Regions 2, 11, 17, 18, and 29 exhibited low detection in one or more datasets across genotypes and were therefore excluded from clustering analyses in all genotypes. Clustering was performed using the Louvain algorithm, with the resolution parameter optimized through trial and error across four values (0.5, 0.8, 1.0, and 1.2). Resolution 0.8 was selected for its effectiveness in generating distinct clusters without over-clustering, which would generate structures that appear largely identical by eye. t-SNE was employed for visualizations; however, the clusters did not appear as completely distinct entities due to the inherent self-similarity of the chromosome data. For the down sampled data used in the time series analysis, a resolution of 1.0 was chosen due to the smaller number of traces per embryonic stage. Clusters for each dataset were ordered and displayed according to boundary strength. Co-clusters were ranked by the number of traces they contained.

#### Median matrix comparison

Matrices from different genotypes were subtracted from one another. To test for significance, we performed Welch’s t-test on all pairwise distance measurements for each interaction. This one-tailed t-test assessed whether the mean of wild type chromosomes is significantly less than the mean of the other sample, assuming that the two samples had unequal variances.

#### Down sample the chromosome between genotypes

To compare chromosome traces across three different genotypes (wild type, H3K9me mutant, and Condensin mutant), we implemented the following steps to mitigate any potential effects of trace count and age distribution. First, the ages associated with each chromosome were binned into 5-age intervals to facilitate age-grouped comparisons. We identified unique binned age groups for each dataset, retaining only the age bins common to all three conditions. For each shared age bin, we determined the minimum number of chromosomes across the datasets, using this value to guide a random down sampling process that ensured equal representation across conditions. Chromosomes were randomly selected from each dataset to match the minimum count within each age bin. This down sampling allowed for age-matched comparisons across all experimental conditions, effectively eliminating potential confounding factors related to differences in age distribution. We note that the matrices remained unchanged after down sampling.

#### Iterative Down sampling and Wilcoxon Testing of Spatial Distances

To assess differences in spatial distance distributions between conditions, we performed multiple random down sampling with repeated statistical testing. For each dataset, we randomly selected n=1000 distances per iteration, repeated across 100 independent iterations. In each iteration, we computed the median distance and tested pairwise differences between datasets using the Wilcoxon rank-sum test (differences in medians).

For each pairwise comparison, we quantified the fraction of iterations yielding p<0.05 (denoted as Significance), which reflects the robustness of the observed difference across random subsets of the data.

#### Radius of Gyration (RoG)

The Radius of Gyration (RoG) was calculated for chromosome traces containing a minimum of 25 regions, as described in Sawh et al. (2020) and Cheng et al. (2021)^36,47^. To assess significant differences between genotypes, we performed a t-test to compare the RoG values across the different groups.

#### Cooperative Chromatin Interactions

To evaluate cooperative interactions among genomic regions, we calculated the probabilities of these regions exhibiting such interactions in comparison to the probabilities of unconditional contact as in Bintu et al. (2018) and Cheng et al. (2021) ^47,52^. We employed a distance threshold of 500 nm, which was selected as it closely approximates the mean distance between adjacent regions.

For each triplet of segments (A, B, and C), we defined the unconditional probability of contact between segments B and C as the percentage of traces exhibiting B-C contact among all traces in which both B and C were detected. Subsequently, we calculated the conditional probability of B-C contact given the occurrence of A-B contact. This was done by first identifying all traces with A-B contact and computing the probability of B-C contact within this subset. Conversely, the conditional probability of B-C contact given the absence of A-B contact was defined as the percentage of traces showing B-C contact among those that did not exhibit A-B contact.

#### Co-clustering and Chi-square test for co-clusters

For co-clustering analysis, the N2, Condensin(-), and H3K9me(-) datasets were extracted from the down-sampled data and clustered together following the method described above in the Clustering section. Chi-square tests for independence were conducted to assess differences in the distribution of chromosome types across clusters. The Chi-square statistic was computed using the approach described in Gutnik et al. (2024) ^41^. Cramér’s V was calculated to measure the strength of the association between chromosome types and clusters while accounting for the influence of sample size.

To identify which cluster and chromosome type combinations contributed most to the Chi-square statistic, standardized residuals were calculated. The standardized residuals quantified the deviation of the observed frequency from the expected frequency under the assumption of independence, providing insight into specific interactions driving the observed associations.

#### Power law fitting

To assess the significance of differences in scaling parameters between the three datasets, we first performed log-log linear regressions to estimate the scaling exponent s and step sizes a at genomic distances of 1 Mb and 100 kb. For each dataset, the regression model was log(D)=log(a)+slog(G) where D is the median spatial distance and G is the genomic distance. Confidence intervals for regression coefficients were used to calculate standard errors, which were propagated to estimate uncertainties in step sizes at 1 Mb and 100 kb, the latter computed as A= a × (0.1) To compare the parameters between datasets, pairwise Z-tests were performed using the estimated parameters and their standard errors:

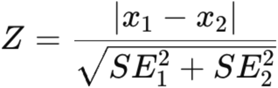

where x represents either the step size or the scaling exponent, and SE their standard errors. P-values were computed from the Z-scores, and to account for multiple comparisons, a Bonferroni correction was applied by multiplying each p-value by the total number of tests performed. Differences with Bonferroni-corrected p-values below 0.05 were considered statistically significant.

### Polymer Modeling

#### Multivariate Normal (MVN) Sampling

To model chromatin folding, we used a Gaussian framework in which polymer configurations are drawn from a multivariate normal (MVN) distribution. Pairwise interactions between loci are encoded in an interaction matrix Λ, which specifies the covariance matrix Σ = Λ^-1^ that fully constrains the MVN distribution. Specifically, for a polymer with *n* monomers, configurations are sampled from the distribution:

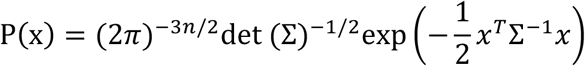

In the interaction matrix Λ, attractive interactions were assigned negative values, while repulsive interactions were positive. The interaction matrix was constructed as a Kirchhoff matrix by setting each diagonal entry such that its corresponding row sums to zero. Because the Kirchhoff interaction matrix has zero determinant due to translational symmetry, we removed the last row and column and fixed the corresponding monomer at the origin (0,0,0). This eliminates the global translational degree of freedom and ensures the matrix can be used for MVN sampling. To generate three-dimensional structures, the interaction matrix was replicated across each of the three diagonal blocks of the full precision matrix. Chromatin conformations were sampled from an MVN distribution using PyTorch’s *torch.distributions.MultivariateNormal*, with the interaction matrix provided as the precision matrix.

In both the Free Polymer and Uniformly Attractive Polymer (UAP) models, five additional monomers were appended to the 5’ end to eliminate terminus effect, as only the 3ʹ end corresponds to the actual chromatin terminus. These extra monomers were excluded from the final output to ensure alignment with the experimental dataset.

#### Specific setting of each model

In the Free Polymer model, we only included uniform nearest-neighbor interactions. The value of the interaction coefficient is set to 45.6 µm^−2^ so that the mean radius of gyration of the free polymer configurations matches the WT experiment dataset at 0.55 µm. Pairs with longer distance are set to zero interaction strength (Fig S8F).

In the UAP model, we set interaction strength based on the distance between the pair of regions. The values are adjusted to fit the folding relationship (Fig S8E). Four different values are used: 7.098 µm^−2^ for nearest neighbors (0.1 Mb), 0.098 µm^−2^ for pairs 0.2–0.9 Mb apart, 0.049 µm^−2^ for pairs 1–1.9 Mb apart, and 0.0147 µm^−2^ for pairs ≥2 Mb apart (Fig S8G). We note that these coefficients are not uniquely determined, as different sets of values might similarly reproduce the observed distance relationship.

In the domain formation models, we specified locations where nearest-neighbor interactions were enhanced and fixed their ratio relative to other nearest-neighbor interactions. These interaction strengths were then optimized using the same procedure as in the UAP model. For models with two compact domains, the same interaction values were used, with only the positions of the enhanced interactions shifted.

## Data and Code availability

Code used in this study was modified from Sawh 2020 or developed by the first author. The data has been deposited in https://github.com/dcpb0/Chromosome-domains.git

## Author Contributions

D.C.P.B. conceived the project, designed the methodology, performed chromosome tracing and data analysis, and wrote the manuscript with input from all authors. J.E.Y. contributed to modeling conceptualization, performed modeling and its analysis, and participated in manuscript review. F.X. designed the probes. T.V. refined analysis scripts. A.N.S. improved the imaging and segmentation pipeline and reviewed the manuscript. P.K. and L.G. contributed to modeling conceptualization. N.M. and D.B. contributed to modeling conceptualization, provided supervision, and reviewed the manuscript. S.E.M. conceived the project, provided supervision and funding, and contributed to writing and review.

## Acknowledgments

We thank the Biozentrum Imaging Core Facility (IMCF) for technical support, and Peter Meister for generously sharing unpublished data and valuable input on the manuscript. We also acknowledge the Caenorhabditis Genetics Center (CGC) for supplying the strains used in this study. This work was supported by funding to S.E.M. from the Biozentrum, University of Basel, SNF 310030_197713 and SNF 320030-227954.

## Supplementary Figures

**Figure S1. Single-molecule tracing replicates show high correlation among themselves and with hi-c data A)** Schematic of the Chromosome V. Region of interest (ROI) traced in this study (dark green). Gray ticks mark Hi-C–defined TAD boundaries ^23^, and the H3K9me3 ChIP–seq signal is plotted below^39^. **B)** Pearson correlation between replicates for wild-type chromosome tracing experiments. **C)** Pearson correlation between median distances obtained from single-molecule tracing and inverse contact frequencies (1/Hi-C contact frequency) for the same region of interest. **D)** Insulation profiles for all clusters

**Figure S2. Condensin I loss affects chromosome V pairwise distances A)** Schematic representation of the experiment using the dpy-28(s939) mutant. **B)** Pearson correlation between replicates for Condensin(-) chromosome tracing experiments**. C)** Significant changes in pairwise distances between wild-type and Condensin(-) mutants, determined by Welch’s t-test. **D)** t-SNE plot of Chromosome V (chrV) clusters in Condensin(-) mutants, where each dot represents a single chromosome trace, and each color corresponds to a cluster.

**Figure S3. Pairwise distance, clustering, and structural changes in H3K9me(-) mutants A)** Pearson correlation between replicates for wild-type chromosome tracing experiments, demonstrating data reproducibility. **B)** t-SNE plot for clusters in H3K9me(-) mutants, where each dot represents a single chromosome trace and each color corresponds to a cluster. **C)** Median pairwise distance matrices (μm) for C4, C5, and C6 clusters in H3K9me traces, annotated with insulation scores. Triangles mark TAD boundary positions, with colors indicating boundary strength. **D)** Significant changes in pairwise distances between wild-type and H3K9me(-) mutants, determined by Welch’s t-test**. E**-**F)** Radius of gyration for wild-type, H3K9me(-), and Condensin(-) traces, down sampled to ensure an equal number of measurements across conditions. Statistical significance was determined using a t-test. n indicates the number of traces.

**Figure S4. Median matrices after downsampling show no changes in interaction patterns A)** Age distribution of the downsampled dataset used in Figure 4 (see Methods for details). **B)** Median pairwise distance matrices (μm) after downsampling to match the distribution in (A). **C)** t-SNE plot of the co-clustering analysis (as in Fig. 4A), with each dot representing a single chromosome trace and colors denoting genotypes. **D)** t-SNE plot of clusters from the co-clustering analysis; dots represent single traces, colored by cluster identity. **E)** Median pairwise distance matrices (μm) for the clusters in (D), before splitting by genotype. **F)** Median pairwise distance matrices (μm) for cluster 6.

**Figure S5. Developmental progression of chromatin structure across later embryonic stages A)** Median pairwise distance matrices (μm) for co-clusters from all genotypes combined at the 2-to 4-cell stage, as shown in Figure 5A. **B)** Standardized chi-square residuals from the co-clustering analysis in (A). High positive values indicate enrichment of traces from a genotype within a cluster, while negative values indicate depletion. **C-E)** Median pairwise distance matrices (µm) for co-clusters from embryos at later developmental stages: 9–40 cells (C), 41–80 cells (D), and 81–140 cells (E), split by genotype. Left panels show genotype contributions.

**Figure S6. Onset of tad-like structures and heterochromatin formation in early embryos A)** Zoomed-in view of the scaling of mean pairwise distance versus genomic distance below 500 Kb for wild-type (blue), H3K9me(-) (red), and Condensin(-) (green) across the five developmental stages shown in Figure 5E. **B)** Scaling exponent (*s*) and **C–D)** step sizes (*a*) at 1 Mb and 100 kb, estimated from log–log linear regression of spatial vs. genomic distances. Error bars represent propagated standard errors. Pairwise Z-tests with Bonferroni correction were used to assess significance; corrected *p*-values are shown below each plot. **E)** Nuclear Volume Across Developmental Stages and Genotypes (μm^3^).

**Figure S7. Representative single-chromosome spatial distance maps across experimental and simulated conditions A–D)** Randomly selected single-trace spatial distance matrices (μm) for wild-type (A), Condensin(-) (B), free polymer simulations (C), and UAP simulations (D). Each matrix represents a single chromosome trace, highlighting the variability in domain organization within and across conditions.

**Figure S8. Comparison of chromatin structure between experimental data and polymer models A)** Representative single-chromosome spatial distance maps (μm) showing domain-like structures in experimental traces. **B)** Boundary probability profile with blue arrows indicating regions of high boundary probability in wild-type embryos. **C)** Distribution of boundary strength in wild-type and H3K9me(-) embryos. A two-sample Kolmogorov–Smirnov (K–S) test was used to compare the distributions of boundary strength and boundary probability between genotypes. **D)** Median pairwise distance matrices (μm) for simulated chromatin structures using the Free Polymer and Uniformly Attractive Polymer (UAP) models, compared with experimental data from the chromosome V region. **E)** Scaling relationships between genomic distance and mean spatial distance for the experimental data, free polymer, and UAP simulations. **F)** Interaction matrix used in the Free Polymer model; asterisks (*) mark additional monomers included to avoid terminus effects. **G)** Interaction matrix for the Uniformly Attractive Polymer (UAP) model. **H)** Interaction matrices for models with enhanced intra-domain interactions (3×, 10×, 100×). **I)** Median pairwise distance matrices (µm) for simulated chromatin structures in (H). **J)** Boundary probability profiles corresponding to (I). **K)** Interaction matrices for the 100× model with two domains separated by 1, 5, or 9 regions. **L)** Example single-molecule distance matrices for the single-domain models in (H). **M)** Example single-molecule distance matrices for the two-domain models in (K).

**Figure S9. Comparison of simulated polymer models with wild-type experimental structures A)** Co-clustering distribution of Free Polymer simulations with wild-type experimental structures. **B)** Co-clustering distribution of Uniformly Attractive Polymer (UAP) simulations with wild-type experimental structures. **C)** Contribution of Free Polymer and experimental data to the clusters. **D)** Contribution of UAP and experimental data to the clusters.

