## Supplementary figures and images for "Single-molecule chromosome tracing reveals a diversity of megabase heterochromatin domains"

### Supplementary Figures

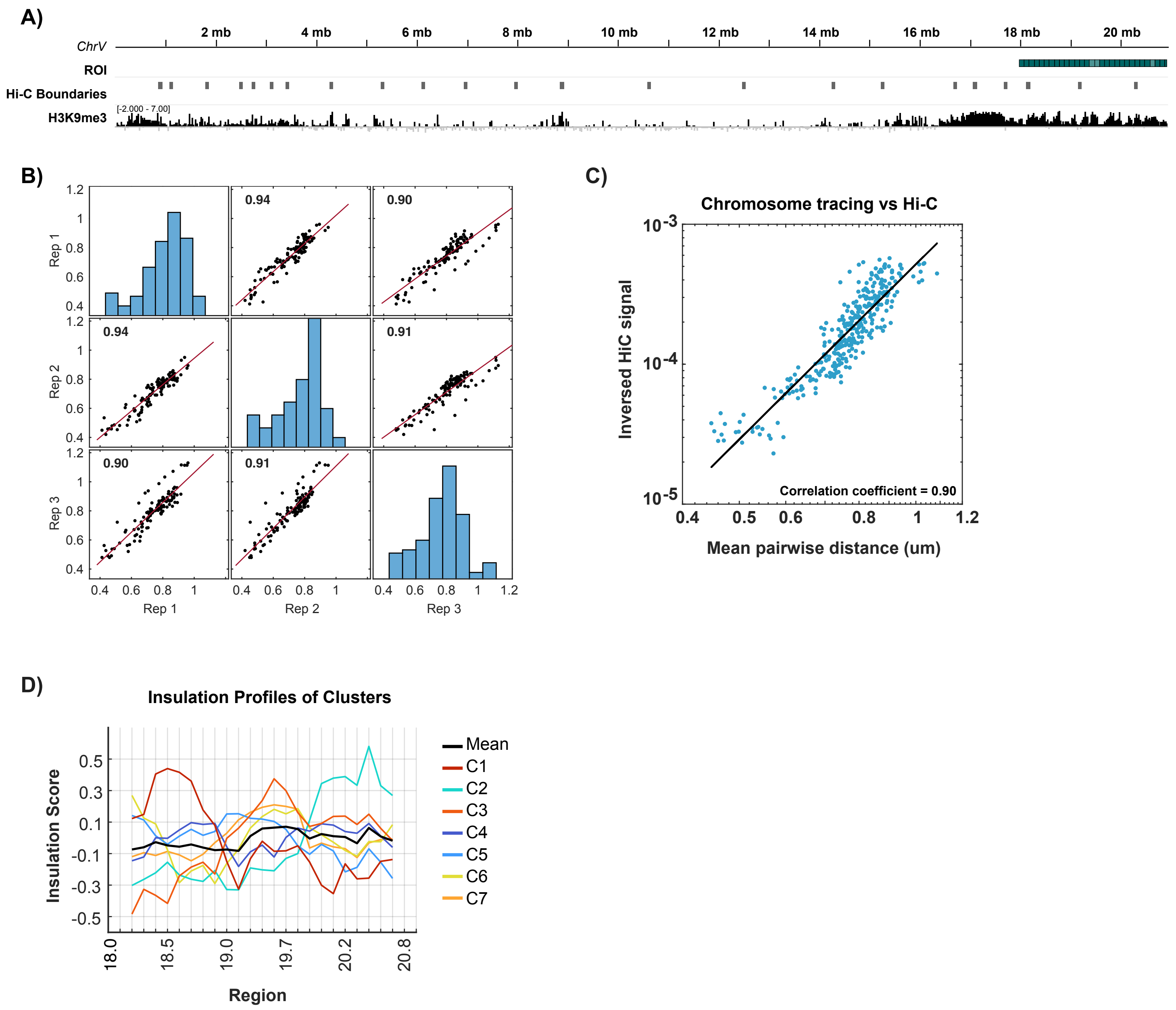

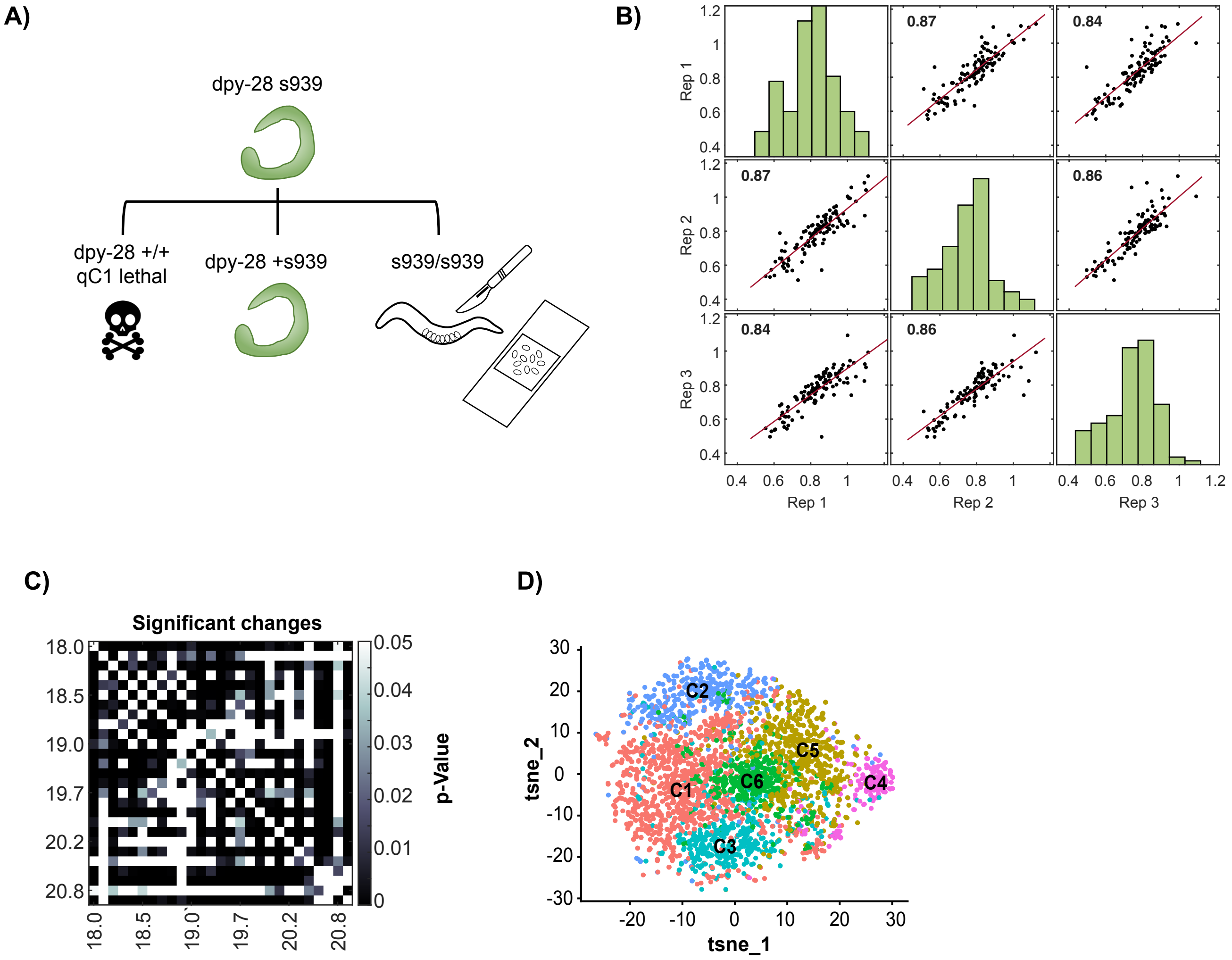

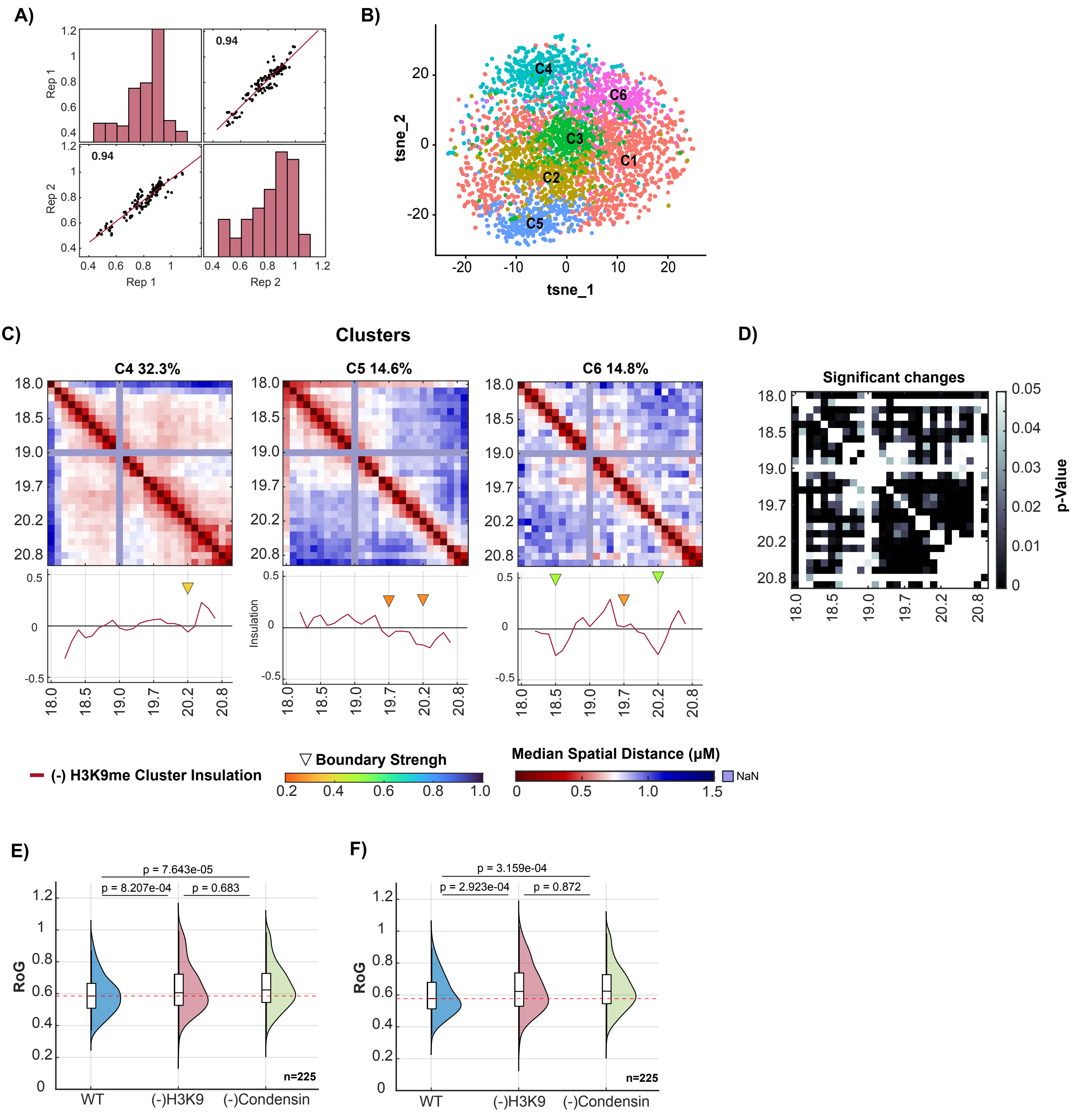

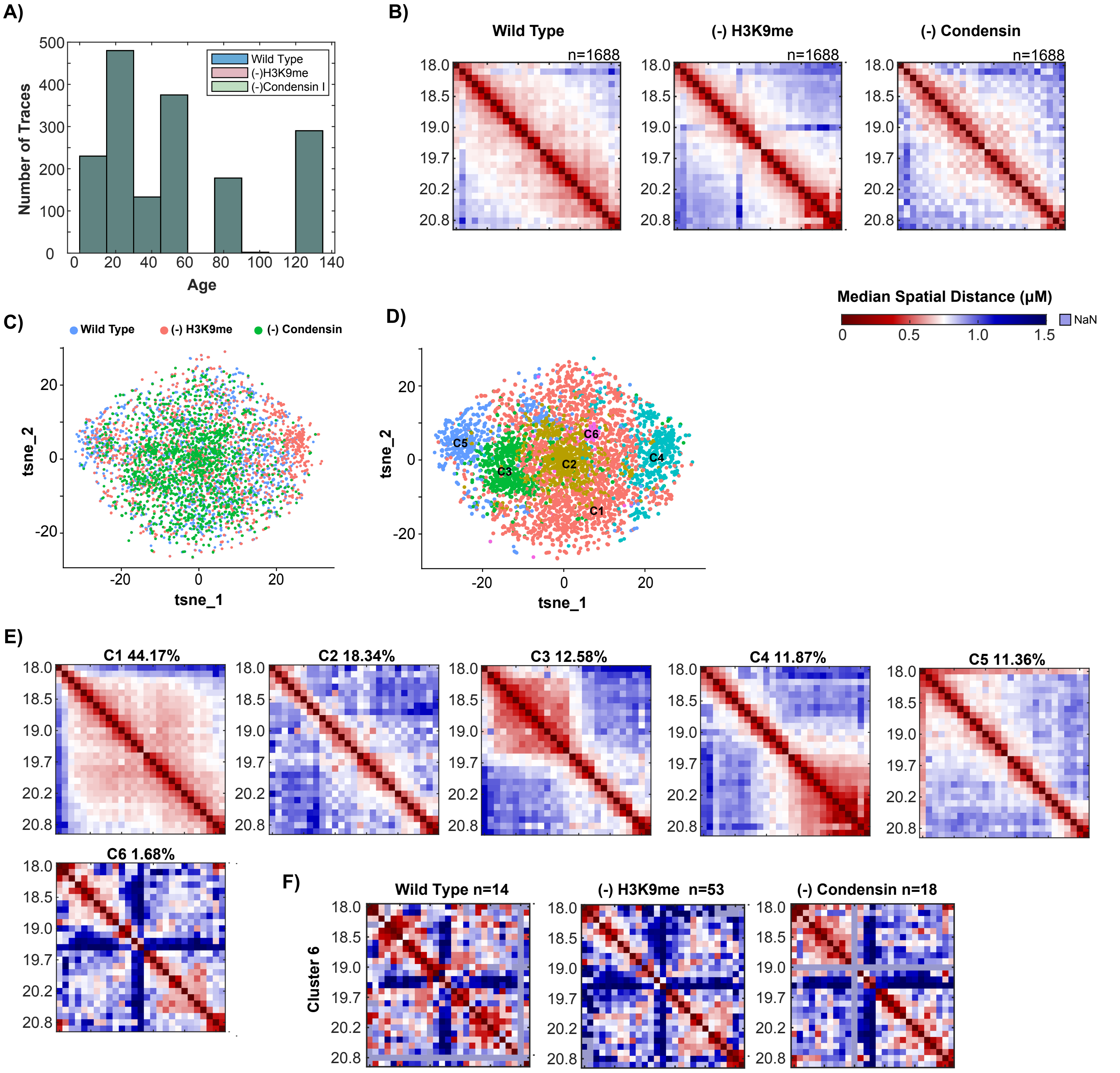

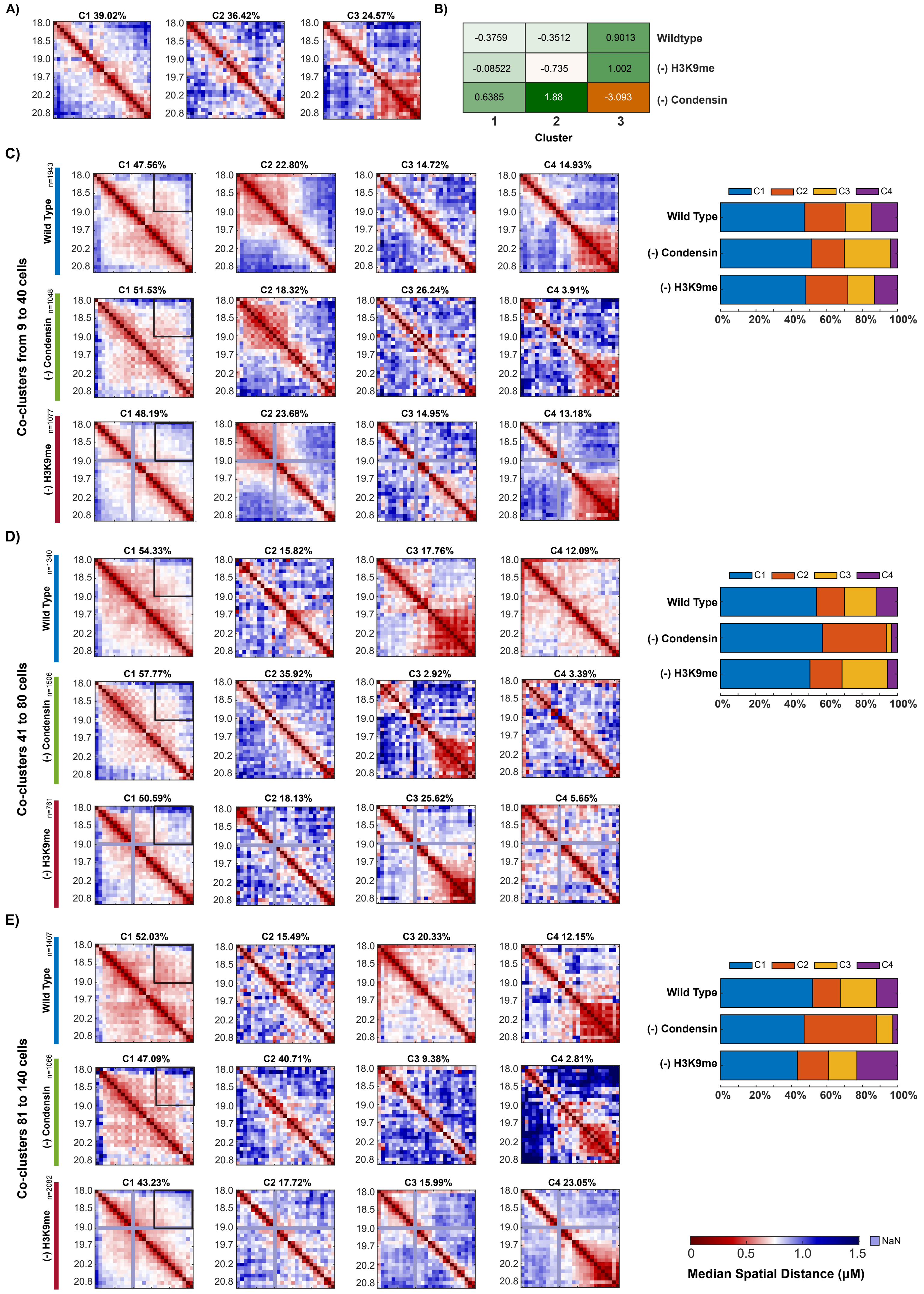

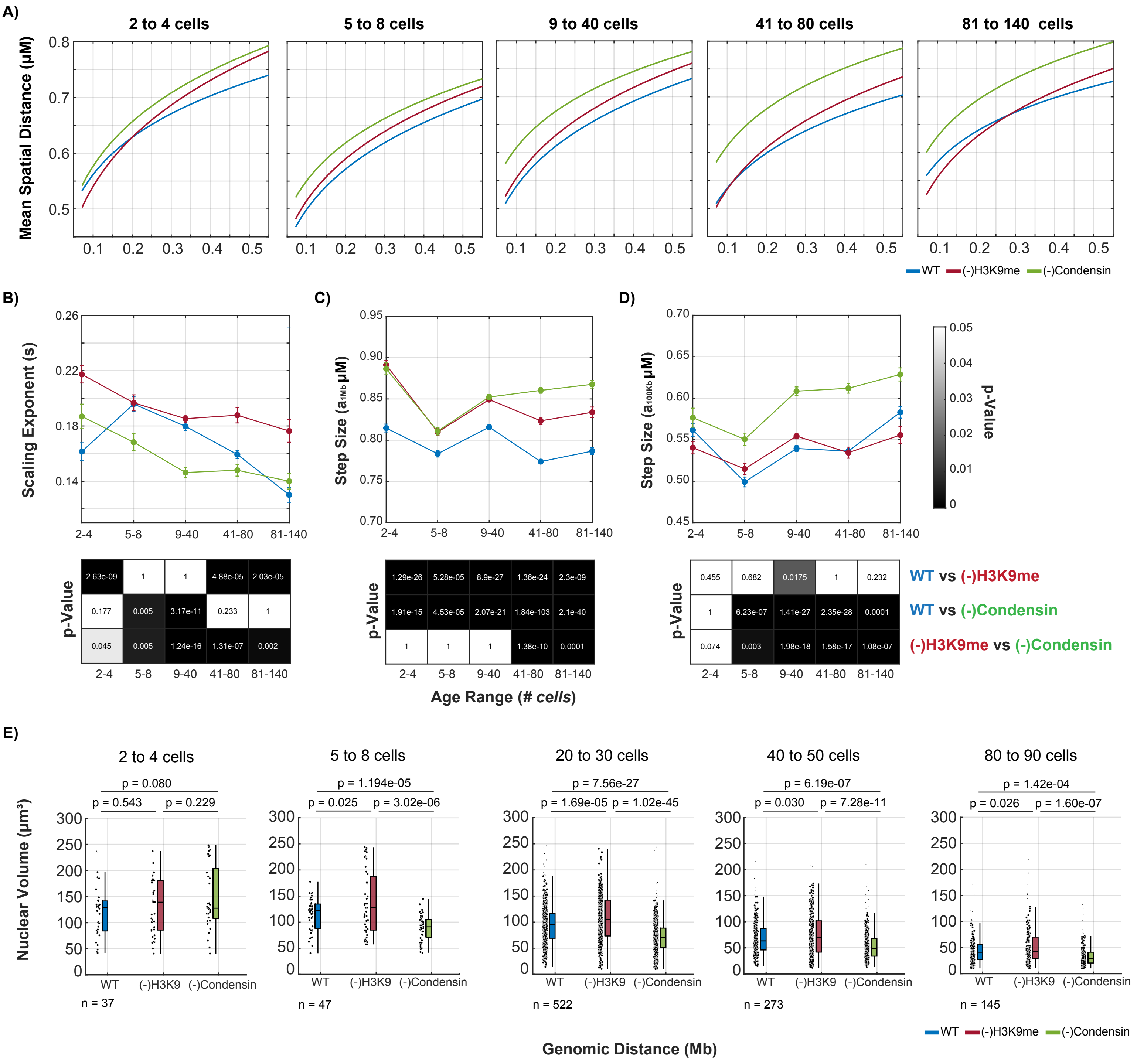

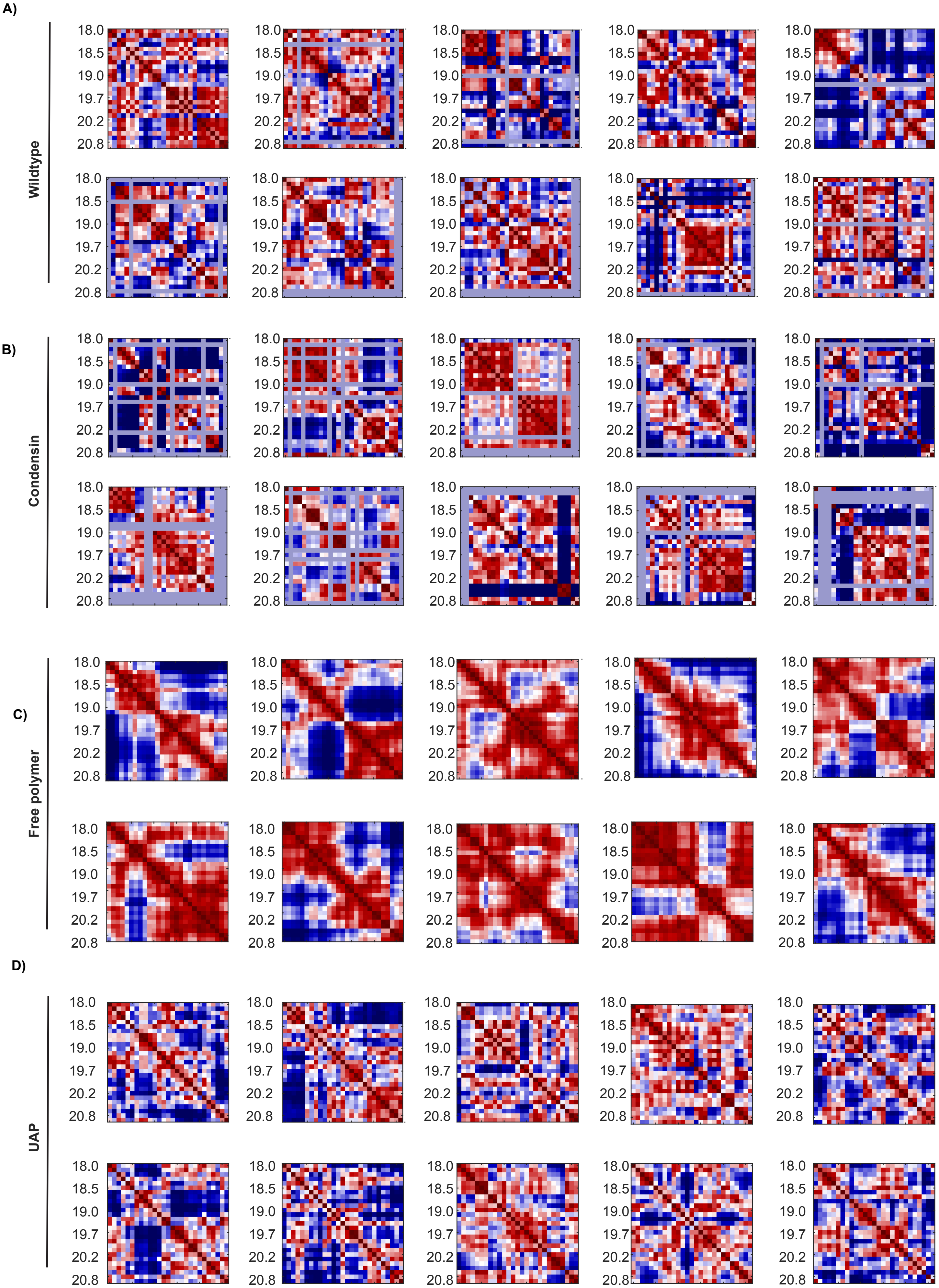

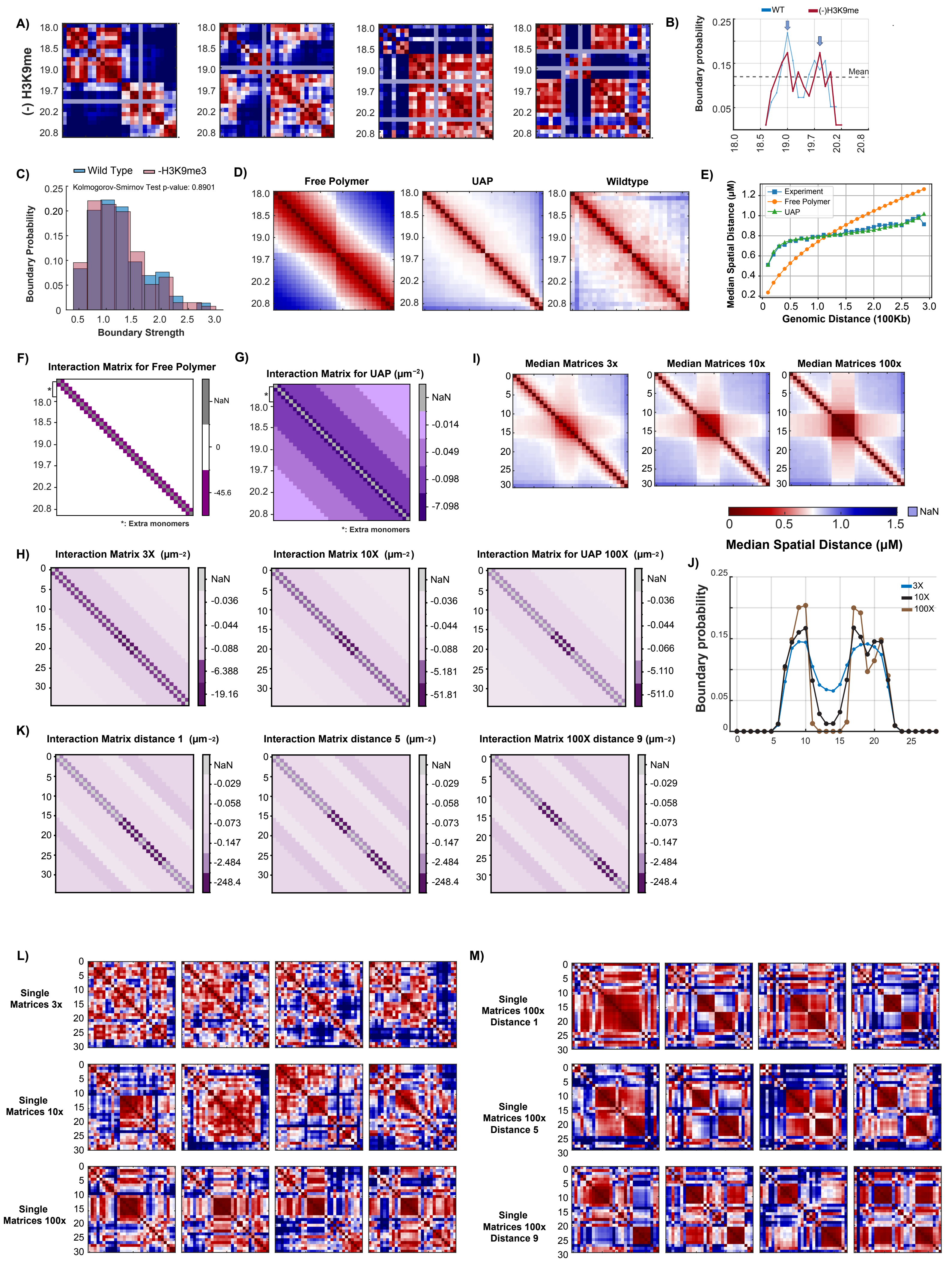

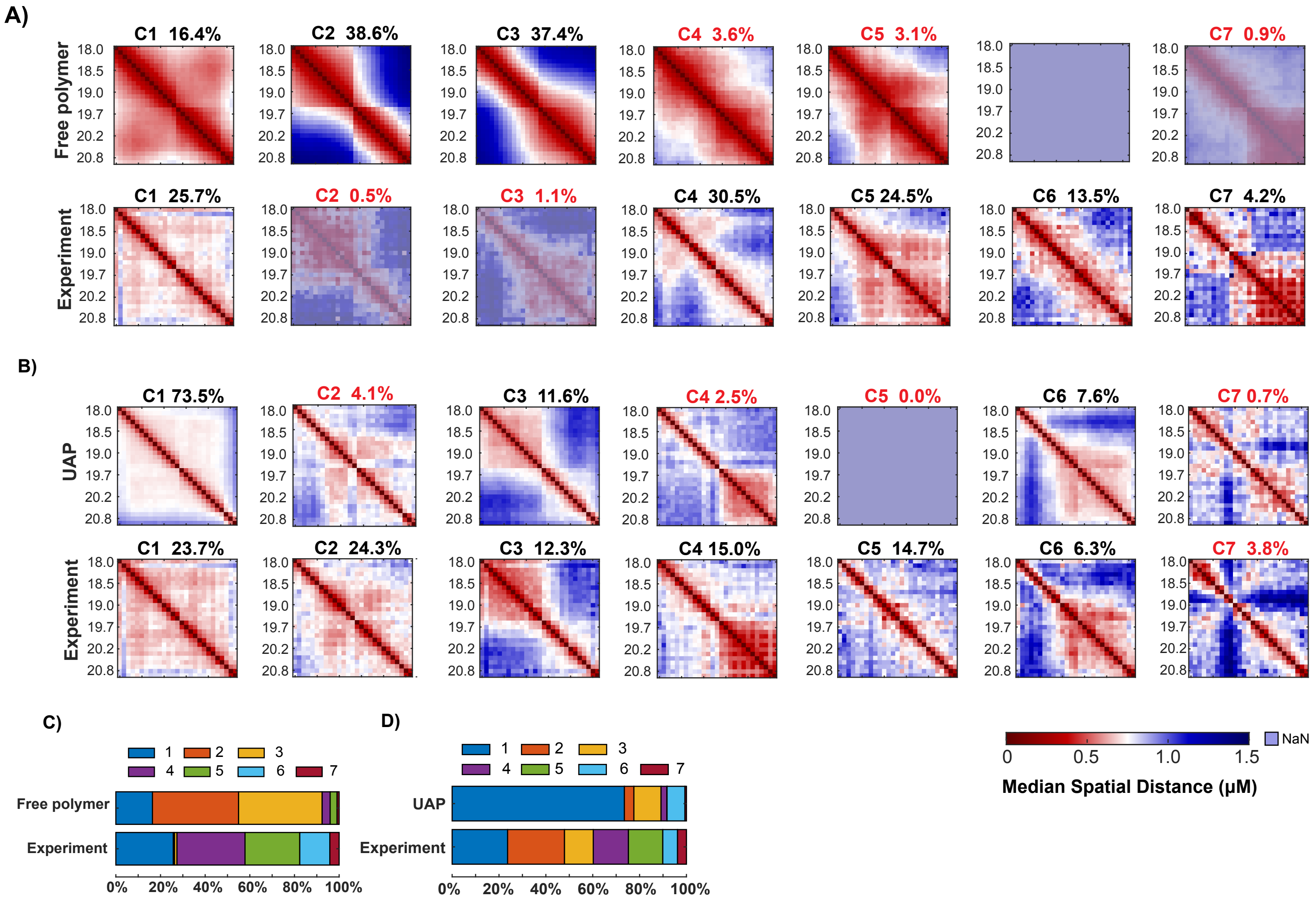
